# A Divide-and-Conquer Method for Scalable Phylogenetic Network Inference from Multi-locus Data

**DOI:** 10.1101/587725

**Authors:** Jiafan Zhu, Xinhao Liu, Huw A. Ogilvie, Luay K. Nakhleh

**Affiliations:** Department of Computer Science, Rice University, Houston, TX 77005, USA; Department of BioSciences, Rice University, Houston, TX 77005, USA

## Abstract

Reticulate evolutionary histories, such as those arising in the presence of hybridization, are best modeled as phylogenetic networks. Recently developed methods allow for statistical inference of phylogenetic networks while also accounting for other processes, such as incomplete lineage sorting (ILS). However, these methods can only handle a small number of loci from a handful of genomes.

In this paper, we introduce a novel two-step method for scalable inference of phylogenetic networks from the sequence alignments of multiple, unlinked loci. The method infers networks on subproblems and then merges them into a network on the full set of taxa. To reduce the number of trinets to infer, we formulate a Hitting Set version of the problem of finding a small number of subsets, and implement a simple heuristic to solve it. We studied their performance, in terms of both running time and accuracy, on simulated as well as on biological data sets. The two-step method accurately infers phylogenetic networks at a scale that is infeasible with existing methods. The results are a significant and promising step towards accurate, large-scale phylogenetic network inference.

We implemented the algorithms in the publicly available software package PhyloNet (https://bioinfocs.rice.edu/PhyloNet).

## 1 Introduction

Phylogenetic networks model non-treelike evolutionary histories, such as those arising when hybridization occurs, and take the shape of a rooted, directed, acyclic graph. Phylogenetic network inference in the genomic era is most often carried out from data obtained from multiple unlinked loci across the genomes of species of interest. To account for the fact that processes such as incomplete lineage sorting (ILS) could co-occur with hybridization, the multispecies network coalescent (MSNC) model was introduced (Yu *et al.*, 2012, 2014) to turn phylogenetic networks into a generative model of gene genealogies, and, subsequently, a wide array of methods for statistical inference of phylogenetic networks under MSNC were introduced (Yu *et al.*, 2014; Yu and Nakhleh, 2015; Wen *et al.*, 2016; Wen and Nakhleh, 2018; Zhang *et al.*, 2018; Zhu *et al.*, 2018; Zhu and Nakhleh, 2018).

Initial evaluations of all these methods on simulated and biological data showed very promising results in terms of the accuracy of the inferences. However, these methods suffer from several major performance bottlenecks. Methods that evaluate the full likelihood (all of the aforementioned methods, except for the pseudo-likelihood method of Yu and Nakhleh (2015)) suffer from the prohibitive computational requirements of likelihood calculations (Zhu and Nakhleh, 2018; Elworth *et al.*, 2018). Currently, computing network likelihood is feasible only for fewer than 10 species and a very small number of reticulations. Second, all the aforementioned methods traverse the space of phylogenetic networks that is much larger than the space of phylogenetic trees, whose size is already exponential in the number of taxa. While the pseudo-likelihood method of Yu and Nakhleh (2015) circumvents the likelihood calculations, albeit in an approximate manner, it does not overcome the problem of exploring the space of the phylogenetic networks. Third, for Bayesian methods, exploring the trans-dimensional space of phylogenetic networks (the number of reticulations changes during the exploration) leads to poor mixing.

In this paper, we propose a method for large-scale phylogenetic network inference that ameliorates all three challenges. The method divides the set of taxa into small, overlapping subsets, builds accurate subnetworks on the subsets, and finally agglomerates the subnetworks into a network on the full set of taxa. By focusing on three-taxon subsets in this paper, the likelihood calculations become very fast, exploring the space of all phylogenetic networks on large numbers of taxa is completely sidestepped. Also, mixing is improved because more iterations of the RJMCMC sampler can be run on three-taxon networks, especially since different subsets can be analyzed independently in parallel. Furthermore, to avoid building all 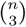 trinets, we provide a Hitting Set formulation of a problem for reducing the number of trinets based on gene trees, and demonstrate that the number of trinets can be reduced significantly without much effect on accuracy.

We implemented our algorithms in PhyloNet (Wen *et al.*, 2018) and studied their accuracy and efficiency. When making use of error-free trinets, we show that the algorithm infers the correct network in all cases, whether making use of all trinets or a significantly reduced subset. When making use of inferred trinets, the algorithm has very good accuracy, where in many cases the correct network is inferred and in all others, a network with small error rate is inferred. This demonstrates the importance of inferring the trinets accurately. Equally important, the method allows for inferring large-scale networks whose inference is infeasible using existing statistical methods.

The closest works to our proposed method here are those of Huber *et al.* (2017); Hejase *et al.* (2018). In (Huber *et al.*, 2017), the authors devised an algorithm that is restricted to combining binet and trinet topologies (no divergence times) into level-1 networks (A phylogenetic network is level-1 if no two cycles in its underlying undirected graphs share a node). The work of Hejase *et al.* (2018) proposed another divide-and-conquer method to infer subnetworks and combine them. However their method makes use of the subnetwork topologies and requires specifying the number of reticulations *a priori*.

The divide-and-conquer method we present here is not only designed to be scalable and make possible the inference of large phylogenetic networks, it also makes use of divergence times so that the estimated network has a time scale. It therefore represents substantial improvement over the previous likelihood-based methods limited in scalability and previous heuristic or summary methods limited in their utility.

## 2 Background

A *phylogenetic network* Ψ on set *𝔛* of taxa is a rooted, directed, acyclic graph (DAG) in which every internal node, except for the root, has in-degree 1 and out-degree 2 (tree node) or in-degree 2 and out-degree 1 (reticulation node). The root has in-degree 0 and out-degree 2, and each leaf has in-degree 1 and out-degree 0. Edges incident into reticulation nodes are the reticulation edges of the network, and all other edges are its tree edges. The leaves of the network are bijectively labeled by the elements of *𝔛*.

For a full probabilistic model, the edges of the network are also associated with continuous parameters as follows. For a given phylogenetic network Ψ, we denote by *V* (Ψ), *E*(Ψ), and *𝔛* (Ψ) the network’s nodes, edges, and leaf labels, respectively. Each edge *b* = (*u, v*) in *E*(Ψ) has a length which is defined by the difference of heights of *u* and *v*, which are denoted by *h*(*u*) and *h*(*v*). Each pair of reticulation edges *e* and *e′* incident into the same reticulation node have inheritance probabilities *γ*_*e*_ and *γ*_*e*_*′* associated with them, which are two non-negative numbers that satisfy *γ*_*e*_ + *γ*_*e*_*′* = 1. Roughly speaking, *γ*_*e*_ denotes the proportion of the genome (in the hybrid population denoted by the relevant reticulation node) that was inherited along edge *e*, and *γ*_*e*_*′* denotes the proportion of the genome that was inherited along edge *e* ′ The network’s topology, branch lengths, and inheritance probabilities fully define the multispecies network coalescent (MSNC) and allows for deriving gene tree probability distributions under ILS and hybridization (Yu *et al.*, 2012, 2014).

For *x* ∈ *𝔛*, we denote by *A*_Ψ_(*x*) and *AR*_Ψ_(*x*) the sets of nodes and reticulation nodes, respectively, on all paths from the leaf labeled by *x*, or node *x*, to the root of Ψ (*AR*_Ψ_(*x*) ⊆ *A*_Ψ_(*x*)). Additionally, we denote *R*(Ψ) to be the set of reticulation nodes in Ψ, with *r*(Ψ) = |*R*(Ψ)|.

*Inference under the MSNC Model.* The data in phylogenomic inferences involves *m* independent loci (genomic regions) consisting of *S* = {*S*_1_, *…, S*_*m*_}, where *S*_*i*_ is the sequence data for locus *i*. Most commonly, *S*_*i*_ could be an alignment of sequences from each of the species under consideration, or *S*_*i*_ is data from a single bi-allelic marker (a vector of 0’s and 1’s), such as a single nucleotide polymorphism (SNP).

The model consists of Ψ, the phylogenetic network (topology and its continuous parameters such as divergence times), and vector Γ of the inheritance probabilities. The likelihood of the model is given by

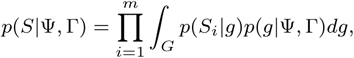

where the integration is taken over all possible gene trees, *p*(*S*_*i*_|*g*) is the probability of the sequence alignment *S*_*i*_ given a particular gene tree *g* (Felsenstein, 1981), and *p*(*g|*Ψ, Γ) is the density of the gene tree (topologies and branch lengths) given the model parameters (Yu *et al.*, 2014). The posterior *p*(Ψ, Γ|*S*) of the model is proportional to

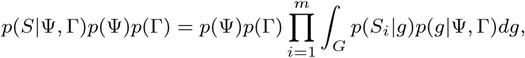

where *p*(Ψ) and *p*(Γ) are the priors on the phylogenetic network (and its parameters) and the inheritance probabilities, respectively.

As discussed above, statistical inference methods under this model suffer from the computational complexity of computing the likelihood, and the challenges with exploring the astronomical and jagged space of phylogenetic networks. Next we describe our method that ameliorates the problem to infer a large network via a two-step approach in which subnetworks are first inferred on smaller data sets of taxa and then the subnetworks are combined to produce the full network.

## 3 Methods

Our divide-and-conquer approach to large-scale phylogenetic network inference on set *𝔛* of taxa takes the following steps:

1. determine a collection of overlapping subsets *𝔛* _1_, *…, 𝔛*_*k*_ of taxa;
2. for each set *𝔛*_*i*_ of taxa, infer an accurate phylogenetic network Ψ_*i*_ (topology, divergence times, and inheritance probabilities) from the sequence data of *𝔛*_*i*_;
3. Combine the *k* subnetworks Ψ_1_, *…*, Ψ_*k*_ into a phylogenetic network on the full set *𝔛* of taxa.

A key issue here is that the sets *𝔛*_*i*_ are small enough so that accurate inference methods, such as (Wen and Nakhleh, 2018), can efficiently and accurately estimate Ψ_*i*_. In this work, we first show the performance when we consider all 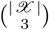 3-taxon subsets, and then propose a technique for reducing this number.

For *Y* ⊆ *𝔛*, we denote by Ψ|_*Y*_ the phylogenetic network restricted to only the leaves labeled by elements of *Y*. We formulate Step (3) in our proposed approach as follows:

- **Input:** Subnetworks Ψ_1_, *…*, Ψ_*k*_ on overlapping sets *𝔛*_1_, *…, 𝔛*_*k*_ of taxa.
- **Output:** Phylogenetic network Ψ with the fewest nodes and edges such that 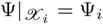 for *i* = 1, *…, k*.

We now describe an iterative algorithm for this problem of combining subnetworks into a full network. The algorithm proceeds in three steps: (1) reconciling and summarizing the node heights across the subnetworks; (2) selecting a starting backbone network (a 3-taxon network in our case) and an order to add taxon-labeled leaves to it; and, (3) iteratively attaching new leaves (*n-*3 of them) according to the computed order until a network on the full set of taxa is obtained.

### 3.1 Reconciling and summarizing the subnetworks

Although two nodes in different subnetworks can correspond to the same node in the true network, a degree of uncertainty is associated with the inferred parameters (mainly their heights) of the two nodes and so they will not exactly match. Those inexact heights will mislead a naïve algorithm that treats differences in heights as strictly pertaining to different nodes, therefore we need to reconcile the parameter estimates in each subnetwork first.

We construct a set 𝒩 of disjoint sets of nodes (each node in each subnetwork has its height). Initially,

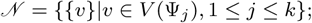

that is, 𝒩 is a set of singletons, one for each node in each of the subnetworks. For every pair (Ψ_*i*_,Ψ_*j*_) of subnetworks, if |*𝔛* (Ψ_*i*_) *∩ 𝔛* (Ψ_*j*_)| *>* 1, we obtain 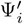 and 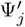 by restricting Ψ_*i*_ and Ψ_*j*_ to *𝔛* (Ψ_*i*_) *∩ 𝔛* (Ψ_*j*_), respectively. By such a restriction, we have two injective mappings from the nodes of 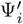 and 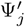 to their corresponding nodes in Ψ_*i*_ and Ψ_*j*_, respectively: 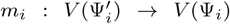 and 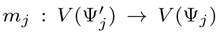. If 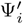 and 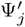 are identical in topology, let 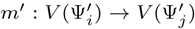 be a bijection between their node-sets. Then for every node 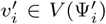, we find the two disjoint sets in 𝒩 containing 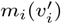 and 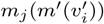, and replace these two sets with their union. If 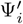 and 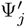 are not identical, we ignore them. In the end, for every node in every disjoint set in 𝒩, we assign the average height of nodes in the same set.

To summarize the height of each node in each subnetwork, here we introduce the “extended height matrix,” or EHM. An EHM *ℳ*_Ψ_ of a network Ψ with *n* leaves is an *n × n* matrix, where element *ℳ*_Ψ_(*x, y*), for taxa *x, y* ∈ *𝔛* (Ψ), is a sorted list of heights of tree nodes in the binet obtained by restricting Ψ to {*x, y*}. We combine 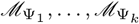 into an EHM *ℳ* for the full network as follows. For *x, y* ∈ *𝔛*, we set *ℳ* (*x, y*) to be the longest list among 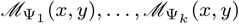. If there are multiple longest lists, the list with smallest lexicographic rank is chosen. For example, if two longest lists (0.1, 0.2, 0.4, 0.9) and (0.1, 0.2, 0.3, 1.0) exist, the latter is chosen. We also define the “pairwise distance sum,” or PDS, for a subnetwork to be the sum of the height of the most recent common ancestor of every pair of taxa in the subnetwork.

### 3.2 Generating a starting network and an order for leaf addition

Here we describe how (1) a starting backbone network is selected, and (2) an order for adding all taxa to it is generated. We assume that a designated taxon *z* has been identified *a priori* to be a member of outgroup with at most 2 members. As this taxon, by definition, is farthest from all ingroup taxa, our task boils down to selecting one of the subnetworks that have *z* as a taxon (when all 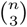 trinets are built, there are 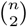 trinets that have *z* as a leaf label). We now describe how to choose one of those as the backbone network.

Let Ψ_*i*_ be a subnetwork whose leaves are labeled by the outgroup taxon *z*, and two other taxa *x* and *y*. We define *s*(Ψ_*i*_) to be 1 if either *x* or *y* is under a reticulation node in any of the *k* subnetworks; otherwise, *s*(Ψ_*i*_) = 0. Furthermore, for two subnetworks Ψ_*i*_ and Ψ_*j*_, we define *d*(Ψ_*i*_, Ψ_*j*_) to be the topological difference (Nakhleh, 2010) of their corresponding restrictions to the set *𝔛* (Ψ_*i*_) *∩ 𝔛* (Ψ_*j*_) of leaves when | *𝔛* (Ψ_*i*_) *∩ 𝔛* (Ψ_*j*_)| *>* 1, otherwise, *d*(Ψ_*i*_, Ψ_*j*_) = 0. We then take as the backbone network the subnetwork

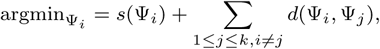

where Ψ_*i*_ iterates over all subnetworks that have *z* as a leaf label, and *k* is the number of subnetworks. If there are multiple subnetworks with the same criterion, the subnetwork with largest PDS is chosen.

Before we add new taxa into the starting backbone, we need to generate an order for attaching new taxa according to the topologies of subnetworks to maximize the correct placement of reticulation nodes. Given two taxa *x, y* ∈ *𝔛* and a collection Ψ_1_, *…*, Ψ_*k*_ of subnetworks, we say that *x* precedes *y*, denoted by *x* 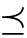 *y*, if 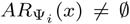 and 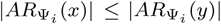 for some Ψ_*i*_. We build a directed graph whose nodes are the taxa set *𝔛*, and edge (*x, y*) is in the graph if and only if *x* 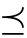 *y*. Then we perform a topological sorting on the directed graph to get an order of attaching missing taxa. Note that there may be cycles in the directed graph; in such a case, when the topological sorting cannot proceed due to a cycle, we break the cycle by removing node *x* (and its incident edges) that appears under a reticulation node in the largest number of subnetworks. The final result is an order of the elements of *𝔛* (minus the three taxa that label the leaves of the backbone network). We create a list of distinct nodes (leaves), each labeled by one taxon, sorted according to the order obtained. The taxa are added to the initial backbone network one at a time according to the computed order. We now describe how each single taxon is added.

### 3.3 Iterative attachment of new taxa

Given the backbone network and the remaining set of taxon-labeled leaves (with their order), we describe how to attach a new taxon to the iteratively growing backbone network. We define the *attachment* of taxon *x* that labels a leaf in subnetwork Ψ_*i*_, denoted by 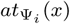, as the set 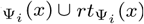, where

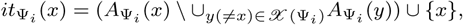

and 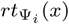 are parent nodes not in 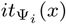 of all nodes in 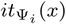. The edges of the attachment, denoted by 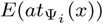, is the set of all edges of Ψ_*i*_ that connect two nodes in the attachment.

We add (leaf labeled by) taxon *x* to the current backbone Ψ_*B*_ as follows. We first compute 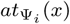 for all *k* subnetworks Ψ_*i*_. Assuming there are *.l* subnetworks that have *x* as a leaf label, we cluster the *.l* attachments by their sizes (all attachments with the same number of nodes in *rt* belong to one cluster), and then choose the single attachment per cluster in which the parent node of the leaf labeled by *x* has the smallest height of all attachments in that cluster. In our implementation, we considered only attachments that have up to 5 nodes in *rt*. Let *H*(*x*) be the set of all resulting attachments (in our implementation, *H*(*x*) contains at most 6 attachments). For each attachment *at*(*x*) = (*it*(*x*) ∪ *rt*(*x*)) ∈ *H*(*x*), we create a set of new backbone networks as follows:

1. For each leaf *x*′∈ *𝔛* (Ψ_*B*_), we generate height-taxon pairs, or HT pairs, according to the overall EHM *ℳ*. The height of the pair is an element of *ℳ* (*x, x*′), and the taxon of the pair is *x*^*′*^.
2. *Resolve* HT pairs by finding the set *P* of positions on the path from *x*^*′*^ (taxon in the pairs) to the root of Ψ_*B*_ where the height of each element in *P* is the height in the pairs. Map the elements of *rt*(*x*) to the positions in *P* in multiple ways. Remove from all the resulting backbone networks any nodes of in-degree 0 except for the original root of the Ψ_*B*_. (Pseudo-code of this step is given in the Supplementary Material.)
3. Remove networks with same topology.

The outcome of this procedure, when applied to all attachments in *H*(*x*), is a set of candidate backbone networks *B*(*x*). We then choose from set *B*(*x*) the network Ψ^*′*^ whose score is minimum. The score of Ψ^*′*^ is defined as follows with respect to each subnetwork Ψ_1_, *…*, Ψ_*k*_:

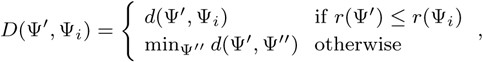

where *d* is the topological distance of Nakhleh (2010) applied to two networks restricted to their shared leaf-set, and Ψ^*′′*^ is taken over all subnetworks of 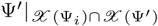 that have *r*(Ψ_*i*_) reticulation nodes.

We choose 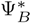 from set *B*(*x*) as the new backbone network on set *𝔛* (Ψ_*B*_) ∪ {*x*} of leaves the network Ψ^*′*^ that minimizes

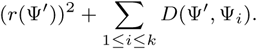

Finally, we reconcile the heights of nodes in 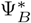 according to subnetworks, by generating a mapping from nodes in 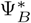 to a set of nodes in the subnetworks, then assign the average of height in each set to the nodes. For inheritance probabilities, we do the same thing for edges in 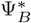.

### 3.4 Asymptotic time complexity

Here we provide a loose analysis of asymptotic time complexity of our merger algorithm if all input subnetworks are trinets. Let the total number of taxa be *n*, and let the total number of reticulations in the true network be *r*. Then it takes at most *O*((*n* + *r*)^2^) to compute the topological difference (Nakhleh, 2010) for two networks which are subnetworks of the true network. Suppose the number of input trinets is *k*. The major time consumption is from the enumeration and evaluation of candidates while attaching new taxa to the growing backbone network.

Suppose we have |*rt*(*x*)| *≤ m* for all attachment in *H*(*x*). For one attachment, there will be at most *O*(*m*! *×* 3^*m*^(*n* + *r*)^*m*^) new backbone networks. In our implementation, we set *m* to 5, which makes the number of candidates *O*((*n* + *r*)^5^). Note that there are far fewer candidates, as demonstrated by our simulation study. A loose upper bound on the time complexity for computing the score for a candidate is *O*(3^*r*^*k*(*n* + *r*)^2^).

The total asymptotic time complexity of our merger algorithm is *O*((*n* + *r*)^5^) *× O*(3^*r*^*k*(*n* + *r*)^2^) *× O*(*k*) = *O*(3^*r*^*k*^2^(*n* + *r*)^7^).

### 3.5 Reducing the number of subproblems

The first step of our method requires inferring a phylogenetic network for every combination of 3 taxa, and this causes the computational complexity of subnetwork inference to be *O*(*n*^3^) given *n* total taxa. If there are 100 taxa, the number of subnetworks to infer will be 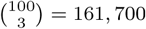, which is an overwhelmingly large number for researchers who do not have access to the largest supercomputers. Therefore, it is important to reduce the number of subnetworks by precomputing which subnetworks are actually needed.

Let *g* be a rooted, binary phylogenetic tree leaf-labeled by set *𝔛* of taxa. For a node *u* in *g*, we denote by *L*(*u*) the set *X*^*′*^ ⊆ *X* that labels the leaves of *g* that are under node *u*. Consider an internal edge *e* = (*u, v*) in *g* (that is, an edge that is not incident with a leaf). Let *v*_1_ and *v*_2_ be the two children of *v*, and let *u*_1_ be the child of *u* that is not *v*. We say that edge *e* is defined by the set {*L*(*v*_1_), *L*(*v*_2_), *L*(*u*_1_)} (that i s, it is a set of three sets of leaf labels). Finally, we say that a triplet of leaf labels {*x*_1_, *x*_2_, *x*_3_} ⊆ *𝔛 covers* edge *e* if

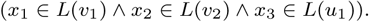

The algorithm we propose for reducing the number of subproblems to solve on a data set of *m* loci is as follows:

1. Let 𝒢 be a set of *m* estimated gene trees, and denote by ℰ (𝒢) the set of all internal edges in the gene trees in 𝒢.
2. Compute a smallest set Δ = {{*x*_1_, *x*_2_, *x*_3_} : {*x*_1_, *x*_2_, *x*_3_} ⊆ *𝔛*} such that each edge *e* ∈ *ℰ* (𝒢) is covered by at least one element of Δ.
3. Infer |Δ| trinets, one for each element of Δ.

We show how computing set Δ can be posed as an instance of the Hitting Set Problem, which allows one to make use of many existing algorithmic developments for this problem. The Hitting Set Problem is defined as follows:

**Input:** A collection *C* of subsets of *S*.

**Output:** Smallest subset *S′* ⊆ *S* that intersects every set in *C*.

To pose our problem of finding a smallest set of 3-taxon subproblems as an instance of the Hitting Set Problem, we define:

- *S* is the set of all 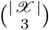 three-taxon subsets of *𝔛*.
- Let edge *e* ∈ *ℰ* (*𝒢*) be defined by the the set {*A, B, C*} of three sets of taxa, as described in the main text. We create set *C*_*e*_ = {{*a, b, c*} : *a* ∈ *A, b* ∈ *B, c* ∈ *C*}. Then,

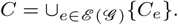

Finding a smallest subset *S′* ⊆ *S* amounts to finding the smallest set of 3-taxon sets on which to infer trinets.

For certain networks (that are automatically identified by the algorithm), the smallest set Δof trinets needs to be enriched with additional trinets that are identified in multiple rounds, a step that we discuss and describe in the Supplementary Material, along with the heuristic we implemented for solving the aforementioned problem.

## 4 Results and Discussion

The way we ran our method is as follows: For each subproblem, MCMC_SEQ (Wen and Nakhleh, 2018) was run and a sample of subnetworks was collected from the posterior. We then selected one subnetwork randomly from the samples of each subset, and applied our merger algorithm. This step was repeated 100 times, and resulted in 100 candidate networks on the full set of taxa. We selected the final network as follows. if a network topology appeared in 50 or more of the 100 networks, it was selected as the final result; otherwise, we identify the most common topology for each of the subnetwork distributions from MCMC_SEQ. Then, we select the network which maximizes the number of subnetworks, contained in that network, which match those topologies. The parameters of the final network are averaged from the networks with same topology.

Since our algorithm for combining subnetworks into a network on the full set of taxa is a heuristic with no established theoretical guarantees, we first set out to study its accuracy on a large number of networks. We then studied the performance of our full approach on simulated multi-locus data sets, and finally analyzed a biological data set.

### 4.1 Accuracy of the merger algorithm

We generated 10,000 16-taxon networks using a birth-hybridization model, and for each network, an outgroup was added to create a 17-taxon network. We restricted each of the 10,000 17-taxa networks to every combination of 3 taxa to produce 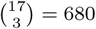 = 680 trinets that were used as input to our merger algorithm that combines the trinets into a network on the full set of taxa. We then inspected the accuracy of the resulting networks. Fig. 1 shows the number of data sets on which the merger algorithm inferred the correct network with 10,000 17-taxon networks. As Fig. 1 shows, in total, 9,838 out of 10,000 inferred networks are identical to their corresponding true networks. When the true network had 0 or 1 reticulations, the algorithm always returned the correct network. Furthermore, the few cases where an incorrect network was returned mostly correspond to large numbers of reticulations (even in those cases, the computed network was very similar to the true one).

**Fig. 1.**
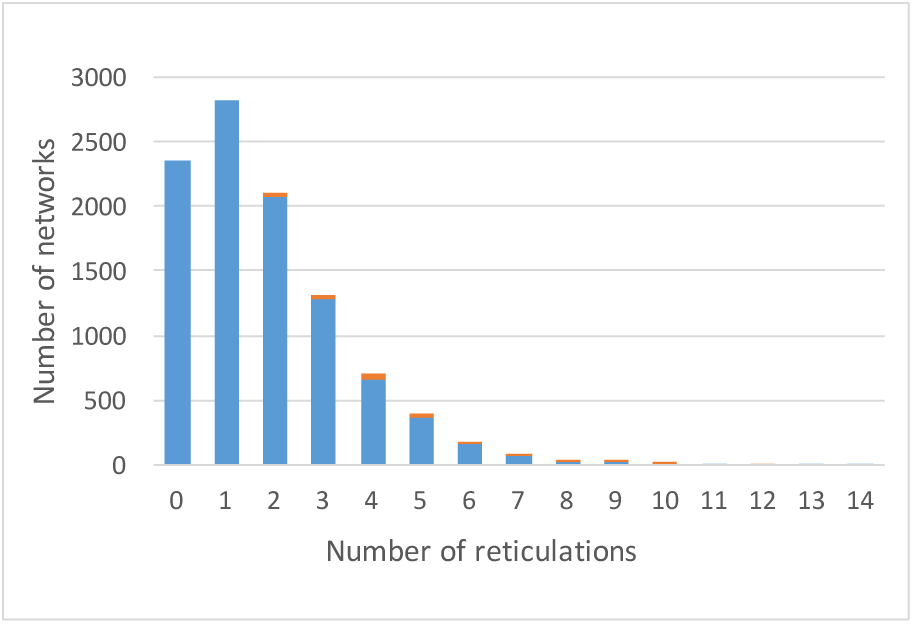
Correctness of inferred networks from correct trinets, categorized by the number of reticulations in the true networks. The numbers of data sets on which the inferred network is identical to or different from the true one are shown in blue and orange, respectively.

To examine the performance of the merger algorithm with and without reduced number of subproblems for large networks, we generated 100 41-taxon networks and 81-taxon networks using a birth-hybridization model (each network had a designated outgroup that did not involve hybridization with any other taxa). We simulated 1,000 gene trees within the branches of each network, using the program ms (Hudson, 2002), and generated the full set of all true trinets as well as subset obtained by our algorithm for reducing the number of trinets. We used each set of trinets as input to our merger algorithm. We inspected the accuracy in terms of whether the inferred network is identical to the true network. The results, as well as other characteristics of the data, are shown in Table 1. When the full set of trinets was used as input, all trinets were inferred in parallel in a single batch. When the reduced set of trinets was used as input, the first batch always consists of the set of reduced trinets being inferred in parallel. However, as we discussed above, in some cases, multiple rounds of enrichment of the reduced set of trinets are performed. Each such round corresponds to an addition batch where all new trinets in that round are inferred in parallel.

**Table 1.**
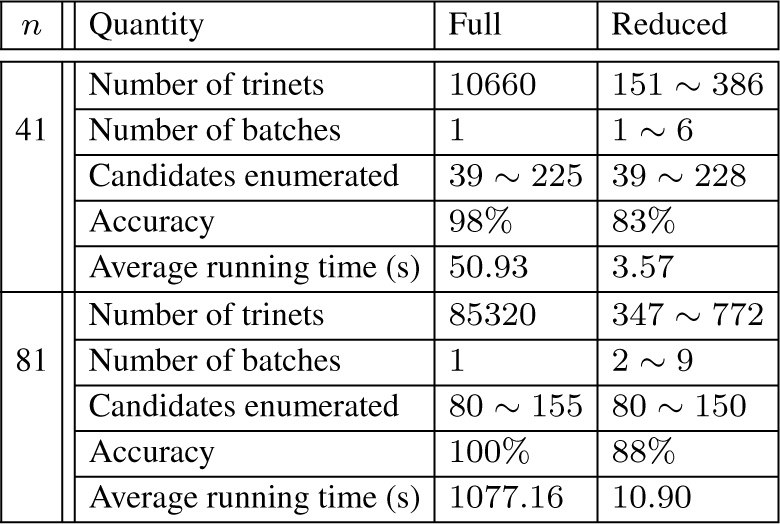
Results of merger algorithm for large networks. Full and Reduced correspond to the full set of trinets and the reduced set of trinets, and *n* is the number of leaves in the network. Each batch consists of multiple trinet inferences that are all run in parallel. ‘Candidates enumerated’ is the number of new backbone networks that are proposed and examined by the algorithm during the full network construction. Accuracy is measured as the percentage of data sets in which the constructed network is identical to the true network. The average running time in seconds is the time it took to construct the full network from the set of trinets.

The table shows several important points. The algorithm achieves almost perfect accuracy on the 41-taxon networks, and perfect accuracy on the 81-taxon networks, when the full set of trinets is used. Our heuristic for reducing the number of trinets achieves two orders of magnitude reduction in the number of trinets, resulting in one or two orders of magnitude reduction in the running time. The accuracy decreases when the reduced set of trinets is used, since some information on the full network is lost by this reduction. We identify the problem of obtaining a better reduced set of trinets as a direction for future research.

One reason the algorithm performs better on the larger networks (81-taxon networks) is that for a fixed number of reticulations, those reticulations would be sparser on a network with 81 taxa than on a network with 41 taxa, making the inference of the former less challenging. Fig. 2 breaks the accuracy results of our algorithm on the 41- and 81-taxon networks by the number of reticulations in these networks.

**Fig. 2.**
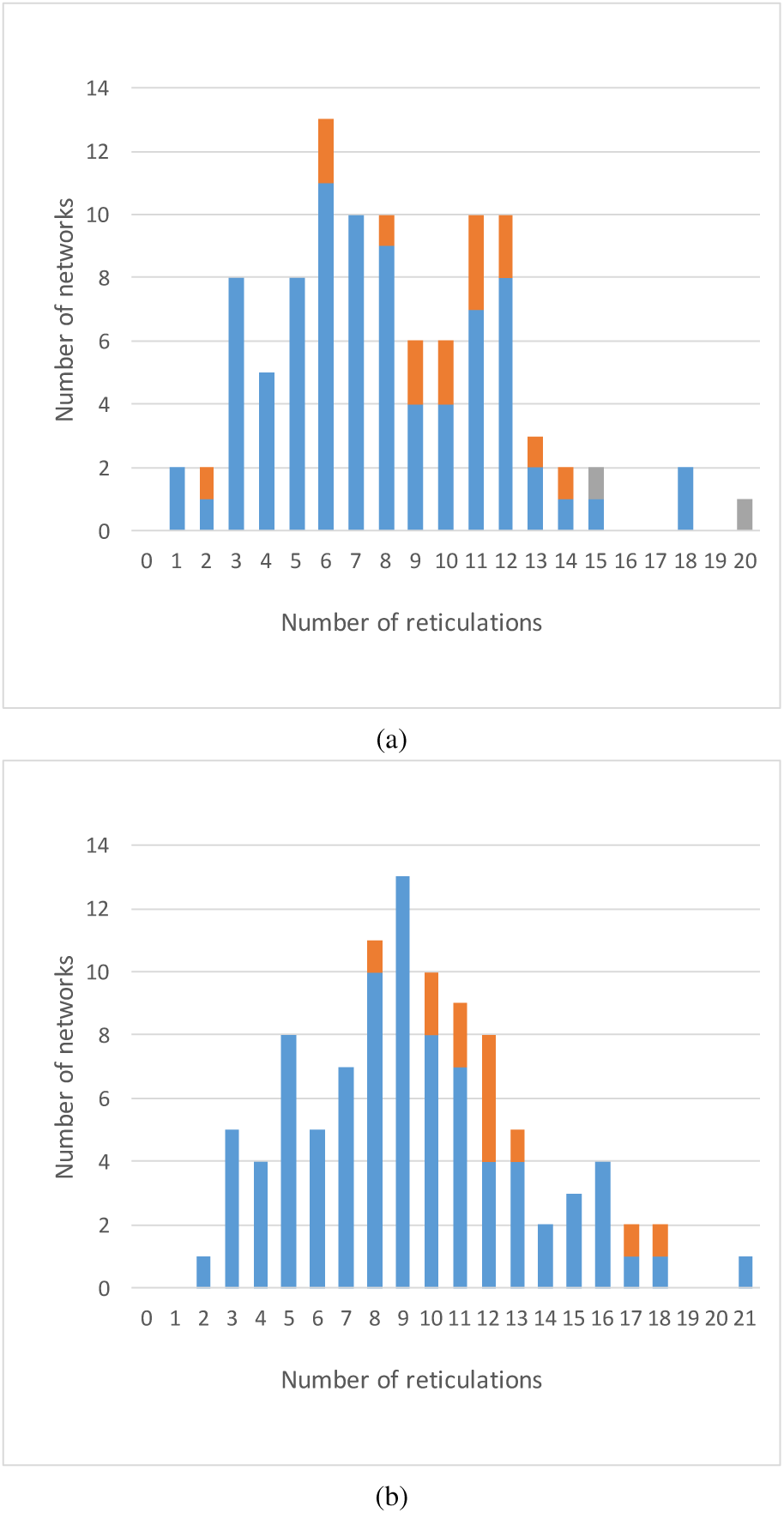
Correctness of inferred networks from correct trinets, categorized by the number of reticulations in the true networks. (a) Results from 100 41-taxon networks. (b) Results from 100 81-taxon networks. Blue: the number of cases where the inferred network is identical to the true one when using either the full or reduced set of trinets. Orange: the number of cases where the inferred network is identical to the true one only when the full, but not reduced, set of trinets is used. Grey: the number of cases where the inferred network is different from the true one, regardless of whether the full or reduced set of trinets was used.

### 4.2 Accuracy on simulated multi-locus data sets

We now set out to study the performance of our approach on simulated multi-locus sequence data, where the method is applied to the sequence data directly. Given that computational complexity of Bayesian inference of trinets (Wen and Nakhleh, 2018), we focus our attention here on a subset of 24 phylogenetic networks that we sampled to reflect varying complexity levels. As discussed in (Zhu *et al.*, 2016; Elworth *et al.*, 2018), the complexity of phylogenetic networks arises not only from the number of leaves or number of reticulation nodes, but also in how the reticulation nodes are structured in the network.To allow for a careful assessment of the accuracy of our approach, we define a simple complexity measure of networks as follows. We define the complexity of Ψ as Σ_*r*∈*R*(Ψ)_ |*L*(*r*)| + |*L*(*p*_1_(*r*))| + |*L*(*p*_2_(*r*))| + | *𝔛 | · |AR*_Ψ_(*r*)|, where *L*(*u*) is the set of leaves under node *u*, and *p*_1_(*u*) and *p*_2_(*u*) are the two parents of reticulation node *u*.

We selected the 24 networks from the 10,000 as follows. All simulated networks with 0 to 5 reticulation nodes were sorted by their complexities. For each of the six numbers of reticulation nodes, we selected four networks: the one with the minimum complexity, the one with the maximum complexity, and the two networks at tertiles. The 24 networks were divided into three groups of 8 “easy” networks (E), 8 “medium-difficulty” networks (M), and 8 “hard” networks (H), and are shown in the Supplementary Material. We used these 24 networks as the ground truth and simulated multi-locus sequence from these 24 networks.

For each of the 24 networks, we generated the full set of all true trinets as well as subset obtained by our algorithm for reducing the number of trinets. Then, for each set of trinets (full or reduced), we perturbed the heights of the nodes in each trinet randomly by 0.1% and repeated this 100 times to obtain 100 “ideal” MCMC-like samples of trinets. We then used the trinet sets as inputs to our merger algorithm and inspected the resulting networks. The algorithm obtained the correct networks in all 24 cases regardless of whether the full or reduced set of trinet “samples” were used. While this result is perfect, Bayesian MCMC in practice is not guaranteed to yield as accurate a sample as the one we used here. Therefore, we next set out to study the performance of the method when we use sequence data of the multiple loci.

For each of the 24 networks, we simulated 100 gene trees, with two individuals per species, for 100 loci using the program ms (Hudson, 2002), and generated sequence alignments of length 1,000 for each locus using Seq-gen (Rambaut and Grassly, 1997) under GTR model. In other words, each locus consists of 34 aligned sequences. For each data set, we inferred subnetworks using MCMC-SEQ (Wen and Nakhleh, 2018) as implemented in PhyloNet (Wen *et al.*, 2018) with 2 × 10^6^ iterations, 1 ×10^6^ burn-in iterations, and one sample collected per 5 ×10^3^ iterations. To obtain the first state for the method, we inferred gene trees for the individual loci using IQ-TREE (Nguyen *et al.*, 2014), optimized their branch lengths using local search, and the resulting gene trees were used as the starting gene trees in the MCMC chain.

For each data set, the running time to infer all trinets is shown in Fig. 3(a). This analysis was performed on NOTS (Night Owls Time-Sharing Service), which is a batch scheduled High-Throughput Computing (HTC) cluster. The average cost to infer all trinets for a data set was 1636.82 CPU-hours, which means it takes about an hour to infer a trinet with a dual-core machine. Since the inferences of trinet are independent of each other, this task is embarrassingly parallel. Fig. 3(b) shows the accuracy of the inferred trinets. The figure shows that the more complex the true network, the harder it is to infer their subnetworks.

**Fig. 3.**
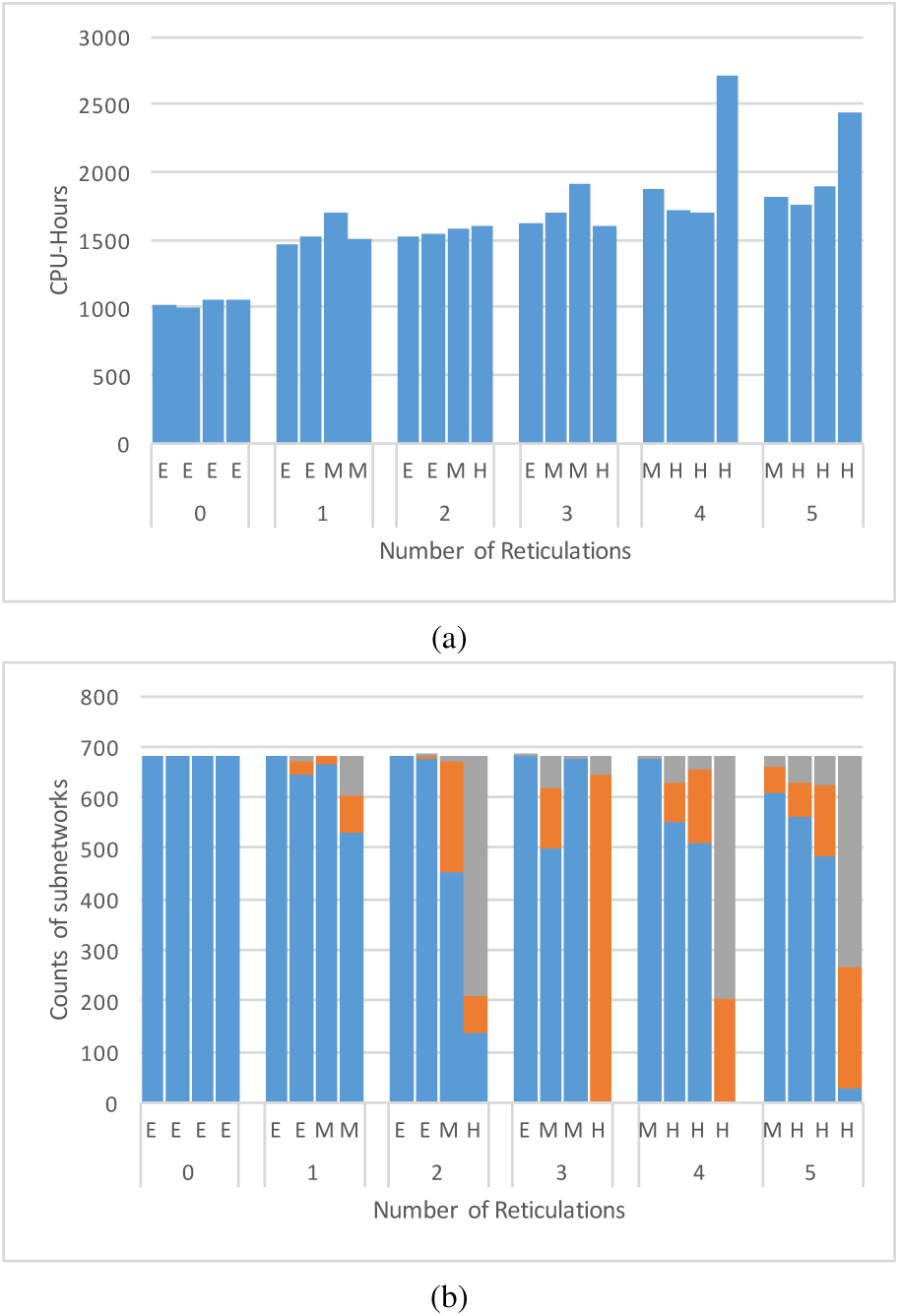
Running times and accuracy for the inferred trinets. (a) The total running time in CPU-hours to infer all trinets for each data set. (b) Accuracy of the inferred trinets. The number of data sets where the inferred trinet is correct (blue), the inferred trinet is inside the true network (orange), and all other cases (grey), are shown.

We then used the inferred trinets as input to our merger algorithm. The merger algorithm ran on a Macbook Pro with 2.9 GHz Intel Core i5. We used both the full and reduced sets of inferred trinets. The reduced sets contains between 61 and 132 trinets, which is a major reduction (especially when considering the running time, as shown in Fig. 3(a)) over the full set, which contains 680 subnetworks. Most data sets only need one batch of inference, 3 data sets need 2 batches, and 1 data set needs 3 batches. The time that our algorithm took to merge the trinets into a full network (repeated 100 times) ranged between 148 and 1538 seconds when the full set of trinets was used, and between 44 and 141 seconds when the reduced set of trinets was used. This shows the additional efficiency gained by reducing the number of trinets.

Finally, we fed the full and reduced sets of trinets to our merger algorithm and compared the inferred networks to the true ones. In measuring the difference between a true network Ψ_*t*_ and and an inferred network Ψ_*i*_, we quantified false positive and false negative rates as follows. We find the backbone 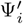 of Ψ_*i*_ and backbone 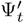 of Ψ_*t*_ whose topological differences (Nakhleh, 2010) are smallest and have the largest number of reticulation nodes among all such pairs of backbones. If the topological difference is 0, the inferred network has a backbone inside the true network. We compute the true positives as the number of nodes remaining in 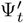, minus the topological difference of 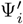 and 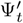. We compute the false positives as the number of nodes deleted from Ψ_*i*_ to 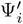, plus the topological difference of 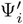 and 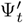. The false negative rate is computed by normalizing the true positives by the number of nodes in Ψ_*t*_ and subtracting it from 1, and the false positive rate is computed by normalizing the false positives by the number of nodes in Ψ_*i*_.

The inferred network was identical to the true network in 12 out of 24 data sets when full set of trinets were used. When the reduced set of trinets was used, 9 inferred networks were identical to their corresponding true networks. We plot the false positives and false negatives for the data sets where the inferred network is not identical to the true one in Fig. 4(a). As the results show, not much accuracy is lost when using the reduced set of trinets. In particular, for four data sets, the false negative rate when using the full set of trinets is higher than its counterpart when using the reduced set. On the other hand, more networks inferred from the reduced set have slightly higher false positive rates. It is important to note here that these results combined with the fact that all 24 inferred networks are completely accurate when using error-free trinets shows that the error in the final networks is mainly due to inaccuracy of the trinets, rather than the merger algorithm.

**Fig. 4.**
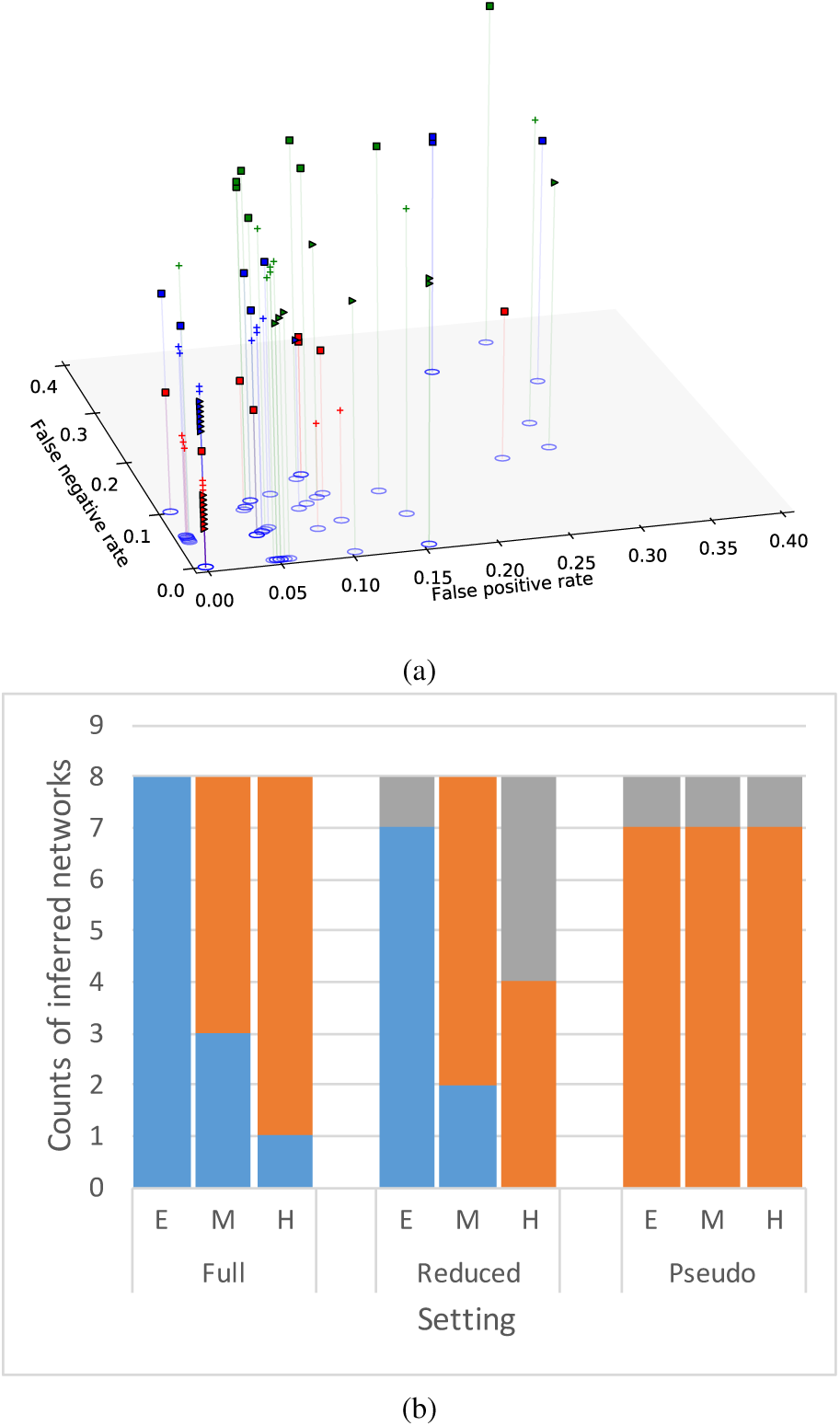
Accuracy of the inferred networks, and comparison to maximum pseudo-likelihood. (a) The false positives and false negatives for the data sets where the inferred network is not identical to the true network. Squares correspond to hard networks, crosses correspond to medium-difficulty networks and triangles correspond to easy networks. Blue, red and green correspond to results based on the full and reduced sets of trinets, and maximum pseudo-likelihood, respectively. (b) The accuracy of our method on the full set of trinets (left set of bars) and on the reduced set of trinets (middle set of bars), and the accuracy of maximum pseudo-likelihood (right set of bars). Blue corresponds to the data sets where the inferred network is identical to the true network; orange corresponds to the data sets where the inferred network contains a backbone network that is present in the true network; grey corresponds to all other cases.

Finally, we compare the accuracy of the method to the only other statistical inference method that can scale to these data sets, namely maximum pseudo-likelihood (Yu and Nakhleh, 2015). As the method of Yu and Nakhleh (2015) requires gene trees as input, we ran it on the gene trees inferred by IQ-TREE, with the maximum number of reticulations set to 5 and the number of runs set to 20. Fig. 4(b) shows the results of this comparison. These results clearly show that our approach here outperforms maximum pseudo-likelihood, and there could be several explanations for this. First, maximum pseudo-likelihood is not good at estimating the correct number of reticulations, so it could be that the networks obtained by the method have unnecessary reticulation nodes. Second, maximum pseudo-likelihood searches the network space and could get stuck in local maxima, whereas our proposed approach here avoids such a search. It is important to also comment on the decreased accuracy of our approach when using a reduced set of trinets. As the set of trinets is much smaller than the full set, the method becomes more sensitive to inaccuracy in the inferred trinets, since when using the full set of trinets, signal from multiple trinets could mask the estimation error. All these results combined show that our proposed approach can produce very accurate results, especially when the individual trinets are accurately estimated.

### 4.3 Inference on an empirical data set

We analyzed a data set of multi-locus sequence alignments of multiple Australian rainbow skinks (Bragg *et al.*, 2018), where 11 taxa with 22 individuals were selected from the full data set. At first we computed the maximum pairwise distance of each locus using IQ-TREE (Nguyen *et al.*, 2014), and we excluded the loci with maximum pairwise distance larger than 0.2, as that would imply impossible deep coalescence times. We then randomly selected 100 loci and used their sequence alignments as the input.

The first step of our method is inferring subnetworks. So we restricted the data set with 11 taxa to every combination of 3 taxa, then we added *Lampropholis guichenoti* into every subproblem to root the subnetworks. Therefore for every subproblem, 4-taxon networks were inferred and the number of subproblems remains 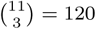. We ran MCMC-SEQ (Wen and Nakhleh, 2018) for 6,000,000 iterations with 3,000,000 burn-in steps, collecting a sample for every 5,000 iterations. We inferred gene trees using IQ-TREE (Nguyen *et al.*, 2014), and their branch lengths were optimized individually using local search. The resulting gene trees were used as the starting point of MCMC chain, and all gene tree topologies were fixed during Bayesian sampling. This analysis was performed on NOTS (Night Owls Time-Sharing Service). We used 2 CPU cores running at 2.6GHz, and 8G RAM for each subproblem. It took 3,670 CPU-hours to infer all subnetworks. Then we used the inferred subnetworks as the input to our merger algorithm to merge them on a Macbook Pro with 2.9 GHz Intel Core i5. It took 53.1 seconds to merge the subnetworks and generate the final result. The inferred network is shown in Fig.5. The ingroup result agrees with the known analysis of this data set. The topological relationships of the *Carlia* clade and the *Lygisaurus* clade are identical to Fig. 2 in (Bragg *et al.*, 2018).

For comparison, we also ran the maximum pseudo-likelihood method of Yu and Nakhleh (2015) on this data set, using the inferred gene trees as the input. The number of runs was set to 10. The number of reticulations allowed was set to 0, 1 and 2. The inferred networks are shown in Fig. 6. The inferred species tree was identical to the backbone tree in the inferred network using our merger algorithm. However, that is no longer the case when reticulations are added by the method.

**Fig. 5.**
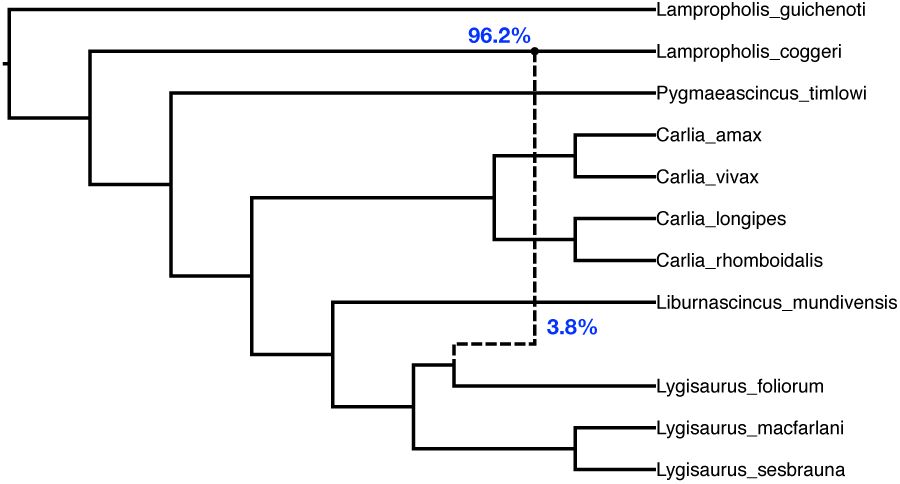
The inferred network for the empirical data set. The reticulation, with inheritance probabilities (blue), is shown by the dashed line.

**Fig. 6.**
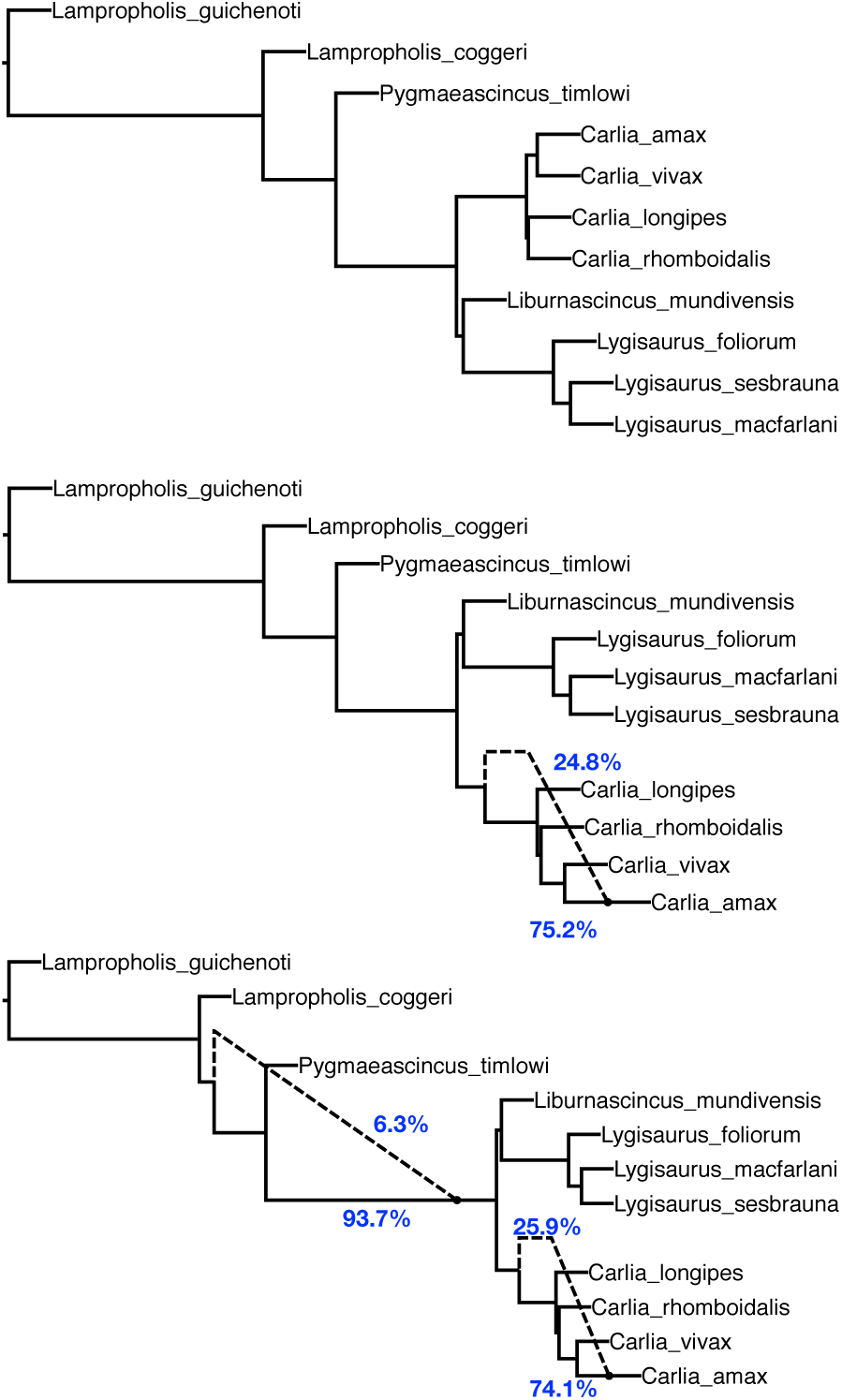
The inferred networks for the empirical data set using maximum pseudo-likelihood. Top: the inferred network when no reticulation was allowed. Middle: the inferred network when 1 reticulation was allowed. Bottom: the inferred network when 2 reticulations were allowed. The reticulations, with inheritance probabilities (blue), are shown by the dashed lines.

## 5 Conclusions and Future Work

In this paper, we proposed a divide-and-conquer approach for large-scale phylogenetic network inference. The approach makes use of inferred subnetworks—topologies and divergence times—on overlapping subsets of the taxa to obtain a phylogenetic network on the full data set. We demonstrated the accuracy and efficiency of our approach on simulated and biological data sets.

While we illustrated the performance of the algorithm on subproblems of size 3 (three taxa), the merger algorithm we introduced works on subnetworks with any number of taxa. There is a tradeoff between the size of the subproblems, the running time, and the accuracy. If the number of taxa in the full data set is *n*, then the full set of subnetworks on *k* leaves consists of 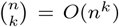. For example, for *n* = 100 and *k* = 5, the algorithm would have to infer on the order of 10^10^ 5-subnetworks. Not only is this number large by itself, but the inference of each 5-subnetwork is much more demanding computationally than that of trinets.

Two bottlenecks of the method are the number of subproblems to analyze, and the time it takes to infer a subnetwork on each subproblem using compute-heavy approaches such as Bayesian inference. To address the former, we introduced a formulation for reducing the number of subproblems to solve and demonstrated its effect on the efficiency and accuracy of the obtained results. However, our solution is a heuristic, and via our reduction of the problem to the Hitting Set Problem, one future direction is to explore the efficiency and accuracy of Hitting Set algorithms. For the latter bottleneck, and while subnetworks can be inferred in parallel on the subproblems, it is important to develop new techniques for accurate estimation of small networks—topologies and divergence times, as these are both used in our approach. Last but not least, while the efficiency of the merger algorithm could be improved, our analyses above show that the two aforementioned bottlenecks are the more important targets for further improvement.

Finally, it is worth mentioning that our merger algorithm makes no assumption on what evolutionary processes were accounted for in the subnetwork inference. In this sense, our merger algorithm can be applied to merge subnetworks inferred under a variety of models (e.g., ILS, gene duplication and loss, and hybridization), as long as the subnetworks’ topologies and divergence times are accurately estimated.

