## Supplementary Materials for "A Divide-and-Conquer Method for Scalable Phylogenetic Network Inference from Multi-locus Data"

### Supplementary Material

#### 1 Algorithm to attach new taxa

Alg. 1 is called to enumerate the ways to connect every attachment $at(x) = it(x) \cup rt(x) \in H(x)$ to current backbone network. To find the positions where $x$ can be connected to, we generate a list, $\mathscr{H}$, of height-taxon pairs, or HT pairs, according to the overall EHM $\mathscr{M}$. Each HT pair $(h', x')$ indicates, there is a tree node at a certain height $h'$ to be the common ancestor of both $x$ and $x'$. A HT pair $(h', x')$ is *resolved*, when there is a tree node in $\Psi'$ which is a common ancestor of both $x$ and $x'$, such that the height of that node is $h'$ within a tolerance.

We copy current backbone network to a draft network $\Psi'$. We modify the draft network by inserting nodes in $rt(x)$ to $\Psi'$, and leaf labeled by $x$ is connected to the draft network and thereby resolves HT pairs. The height of inserted node is set to the height of HT pair. We resolve HT pairs from lowest height. We enumerate the ways to resolve HT pairs by permuting nodes in $rt(x)$, then use depth first search algorithm to search where to connect a node in $rt(x)$.

**Enumerate** (Alg. 1) computes $\mathscr{H}$ as well as calls the depth first search algorithm **EnumerateDFS** (Alg. 2).

**Input**: Network $\Psi_B$. Full EHM $\mathscr{M}$. Taxon $x$. Tolerance $\epsilon$..
**Output**: Set of networks $\Omega$.
$\Omega \leftarrow \emptyset$;
$\mathscr{H} \leftarrow []$;
**foreach** *Taxon* $x' \in \mathscr{X}(\Psi_B)$ **do**
    **foreach** *Height* $h \in \mathscr{M}(x, x')$ **do**
        Append $(h, x')$ to $\mathscr{H}$;
    **end**
**end**
Sort $\mathscr{H}$ according to heights of the entries in ascending order;
Draft network $\Psi' \leftarrow \Psi_B$;
**foreach** $at(x) = (it(x) \cup rt(x)) \in H(x)$ **do**
    **foreach** *Permutation* $r = (r_1, r_2, \cdots)$ *where* $r_1, r_2, \cdots \in rt(x)$ **do**
        Remove the height of $r_1, r_2, \cdots$;
        $\Omega \leftarrow \Omega \cup$ **EnumerateDFS**$(\Psi', at(x), r, \mathscr{H}, x, \epsilon)$;
    **end**
**end**
**return** $\Omega$;

**Algorithm 1**: **Enumerate.**

**Input**: Draft network $\Psi'$. Attachment $at$. List $r$ of $rt$ nodes in $at$. List of HT pairs $\mathscr{H}$. Taxon $x$. Tolerance $\epsilon$.
**Output**: Set of networks $\Omega$.
$\Omega \leftarrow \emptyset$;
`// Clone current draft network and check whether to store it.`
$\Psi'' \leftarrow \Psi'$;
Remove nodes with in-degree 0 in $\Psi''$ until the only node with in-degree 0 is the root;
Remove binary nodes in $\Psi''$;
**if** $\Psi''$ *has no cycle and* $\Psi''$ *has no negative branch length* **then** $\Omega \leftarrow \Omega \cup \{\Psi''\}$ `// If no more nodes to connect, no more search.`
**if** $r$ *is empty* **then return** $\Omega$ $\mathscr{M}_{\Psi'} \leftarrow$ extended height matrix of $\Psi'$;
`// Remove all resolved HT pairs.`
**while** $\mathscr{H}$ *is not empty* **do**
  $(h', t') \leftarrow$ first HT pair in $\mathscr{H}$;
  **if** $h'$ *exists in* $\mathscr{M}_{\Psi'}(x, t')$ *within tolerance* $\epsilon$ **then** Remove $(h', t')$ from $\mathscr{H}$
**end**
`// If all HT pairs are resolved, no more search.`
**if** $\mathscr{H}$ *is empty* **then return** $\Omega$ `// Otherwise resolve the unresolved HT pair with lowest height.`
$(h', t') \leftarrow$ first HT pair in $\mathscr{H}$;
$b \leftarrow$ first node in $r$ and remove first node from $r$;
`// b must have one child according to definition of attachment.`
$b' \leftarrow$ child of $b$;
$h(b) \leftarrow h'$;
**if** $b'$ *is a tree node* **then** $b''_1, b''_2 \leftarrow$ children of $b$ `// Enumerate the ways to resolve current HT pair.`
**for** $k \in \{1, 2, 3\}$ *if* $b'$ *is a tree node else* $k \in \{3\}$ **do**
  **if** $k \leq 2$ **then** Delete edge $(b', b''_k)$ **foreach** $e = (u, v) \in \mathscr{E}(\Psi')$ *and* $e \notin at$ **do**
    **if** $h(u) > h' > h(v)$ **then**
      Break $(u, v)$ into $(u, b)$ and $(b, v)$;
      $\Omega \leftarrow \Omega \cup$ **EnumerateDFS**$(\Psi', at, r, \mathscr{H}, x, \epsilon)$;
      Restore $(u, v)$;
    **end**
  **end**
  **if** $k \leq 2$ **then** Restore edge $(b', b''_k)$
**end**
Restore $b$ in $r$;
**return** $\Omega$;

**Algorithm 2**: **EnumerateDFS.**

#### 2 Reducing the number of subproblems

In the main text, we formulated the problem of reducing the number of trinets to build and posed it as an instance of the Hitting Set Problem. We implemented the following greedy heuristic for solving the problem given an input set $\mathscr{G}$ of gene trees:

- $\Delta \leftarrow \emptyset$;
- **foreach** $e \in \mathscr{E}(\mathscr{G})$
  - **if** $e$ is not covered by an element in $\Delta$
    - Let $\{x_1, x_2, x_3\} \subseteq \mathscr{X}$ be an arbitrary element that covers $e$;
    - $\Delta \leftarrow \Delta \cup \{\{x_1, x_2, x_3\}\}$;
- **return** $\Delta$;

After $\Delta$ is generated by the greedy heuristic, we compute $\Delta'$ by the mapping $S$ from alleles in gene trees to leaves in the full network using another greedy heuristic:

- $\Delta' \leftarrow \emptyset$;
- **foreach** $\{x_1, x_2, x_3\} \in \Delta$
  - $x'_1, x'_2, x'_3 \leftarrow S(x_1), S(x_2), S(x_3)$
  - **if** $|\{x'_1, x'_2, x'_3\}| = 3$
    - $\Delta' \leftarrow \Delta' \cup \{\{x'_1, x'_2, x'_3\}\}$;
  - **else if** $|\{x'_1, x'_2, x'_3\}| = 2$
    - Replace one of repeated species with the outgroup;
    - $\Delta' \leftarrow \Delta' \cup \{\{x'_1, x'_2, x'_3\}\}$;
  - **else if** $|\{x'_1, x'_2, x'_3\}| = 1$
    - Discard;
- **return** $\Delta'$;

To check whether the set of inferred subnetworks is sufficient to be used by the merger algorithm, we use the following algorithm **Enrich** to find additional subnetworks needed. **Enrich** is a naive version of merger algorithm: instead of building a network, it builds a binary tree according $\Psi_1, \cdots, \Psi_{|\Delta'|}$. If it cannot build a binary tree, it provides a set of additional subnetworks needed.

- $\Delta'' \leftarrow \emptyset$;
- Reconcile heights of nodes in $\Psi_1, \cdots, \Psi_{|\Delta'|}$;
- Combine EHMs $\mathscr{M}_{\Psi_1}, \ldots, \mathscr{M}_{\Psi_{|\Delta'|}}$ into $\mathscr{M}$;
- Let $M$ be the height matrix with same dimension as $\mathscr{M}$, and let $M(x, y)$ to be the first item of $\mathscr{M}(x, y)$ if $\mathscr{M}(x, y)$ exists; otherwise $M(x, y) = \infty$;

- Select a starting network $\Psi_t$ from $\Psi_1, \cdots, \Psi_{|\Delta'|}$;
- Let $x_1, x_2, x_3$ be the taxon of $\Psi_t$ such that $M(x_1, x_2) < min(M(x_1, x_3), M(x_2, x_3))$;
- Build a tree $T$ of $x_1, x_2, x_3$, whose internal node is the parent of $x_1$ and $x_2$ with height $M(x_1, x_2)$, and the height of its root is the same as $\Psi_t$;
- Let $Order$ be the order of leaf addition computed from $\Psi_1, \cdots, \Psi_{|\Delta'|}$;
- **foreach** $x$ in $Order$
  - $x' \leftarrow \arg\min_{x'}\{M(x, x')\}$ such that $x'$ is in $T$;
  - **if** $M(x, x') = \infty$
    - **foreach** node $p$ in $T$
      - $y, z \leftarrow$ two arbitrary taxa separated by $p$;
      - $\Delta'' \leftarrow \Delta'' \cup \{\{x, y, z\}\}$;
    - **return** $\Delta''$;
  - **else if** $\exists p$ such that $p$ is an ancestor of $x'$ and $|M(x, x') - h(p)| < \epsilon$
    - **foreach** pair of taxa $y$ and $z$ separated by $p$
      - $\Delta'' \leftarrow \Delta'' \cup \{\{x, y, z\}\}$
    - **return** $\Delta''$;
  - **else**
    - Add a leaf labeled by $x$ to $T$ by creating a common ancestor $p$ of $x$ and $x'$ such that $h(p) = M(x, x')$;
- **return** $\emptyset$;

Finally we can change our first step into **ReducedSubnetworkInference.** (Alg. 3). In this version, the first batch of subnetworks are inferred from $\Delta'$. Then we call **Enrich** to enrich the set of subnetworks iteratively. After running this version of first step, the selected starting network and order of leaf addition from the last call of **Enrich** is kept, and the merger algorithm needs to use the same starting network and order of leaf addition.

**Input**: Multilocus sequence alignments $S_1, \cdots, S_m$.
**Output**: List of subnetworks.
$g_1, \cdots, g_m \leftarrow$ gene tree inference given $S_1, \cdots, S_m$ ;
Get $\Delta'$ using greedy heuristic from $g_1, \cdots, g_m$;
Infer the subnetwork $\Psi_1, \cdots, \Psi_{|\Delta'|}$ with subproblems in $\Delta'$;
$\Delta'' \leftarrow$ **Enrich**$(\Psi_1, \cdots, \Psi_{|\Delta'|})$;
**while** $\Delta'' \neq \emptyset$ **do**
    Infer subnetworks $\Psi_{|\Delta'|+1}, \cdots, \Psi_{|\Delta'|+|\Delta''|}$ corresponding to additional triplets;
    $\Delta' \leftarrow \Delta' \cup \Delta''$;
**end**
**return** $\Psi_1, \cdots, \Psi_{|\Delta'|}$;

**Algorithm 3**: **ReducedSubnetworkInference.**

### 3 True networks in the simulation study

The following is a table of 24 true networks used in our simulation study. The numbers of reticulations, the subjective difficulty rankings are given under networks.

| | |
|---|---|
| 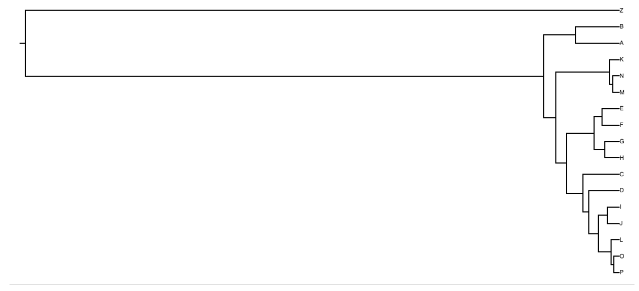 | 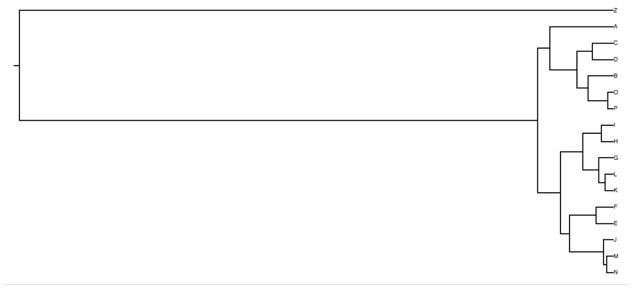 |
| 0 reticulation. Difficulty: Easy. | 0 reticulation. Difficulty: Easy. |
| 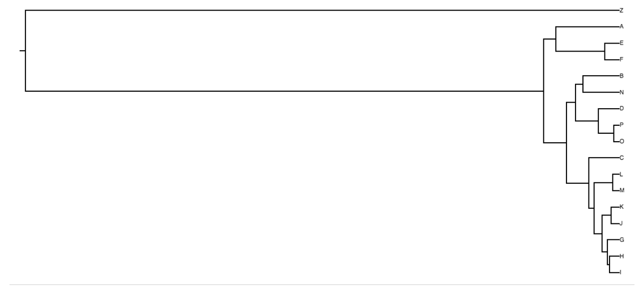 | 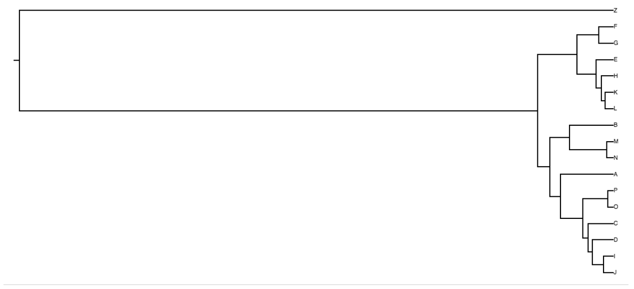 |
| 0 reticulation. Difficulty: Easy. | 0 reticulation. Difficulty: Easy. |
| 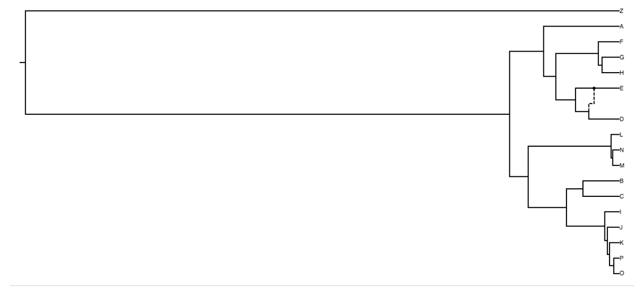 | 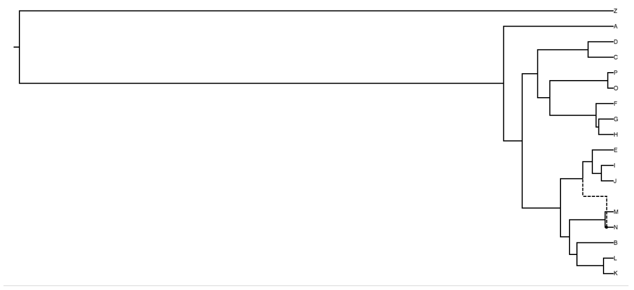 |
| 1 reticulation. Difficulty: Middle. The reticulation is hard to be identified since it only depends on D. | 1 reticulation. Difficulty: Easy. |
| 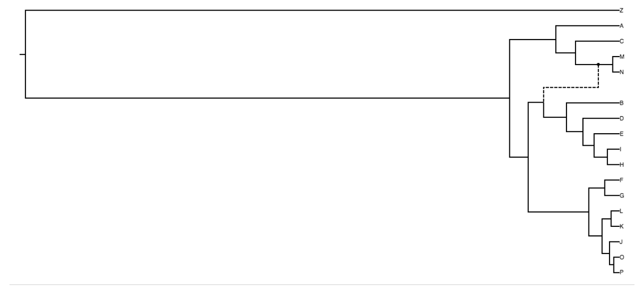 | 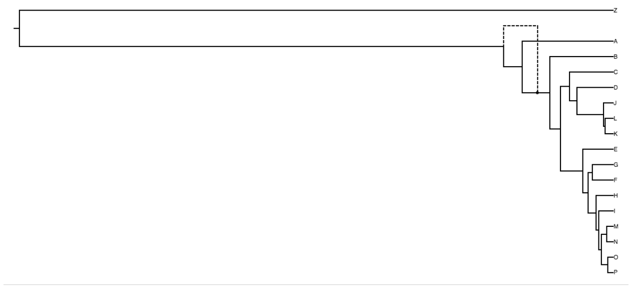 |
| 1 reticulation. Difficulty: Easy. | 1 reticulation. Difficulty: Middle. The reticulation is too deep to be correctly identified. |

| | |
|---|---|
| 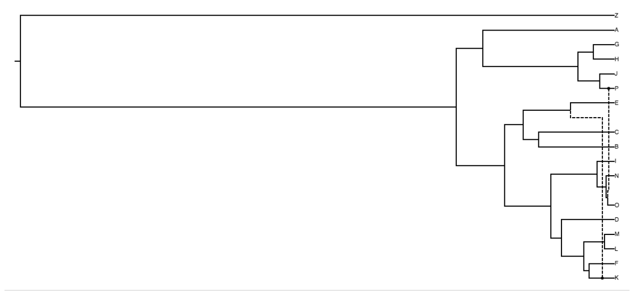 | 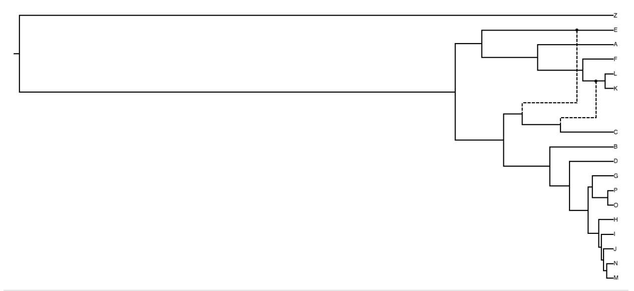 |
| 2 reticulations. Difficulty: Easy. | 2 reticulations. Difficulty: Middle. The dependency of 2 reticulations makes inference not easy. |
| 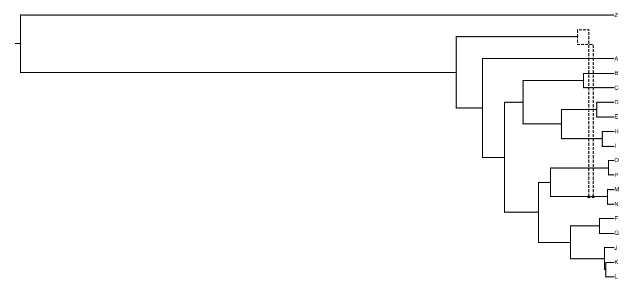 | 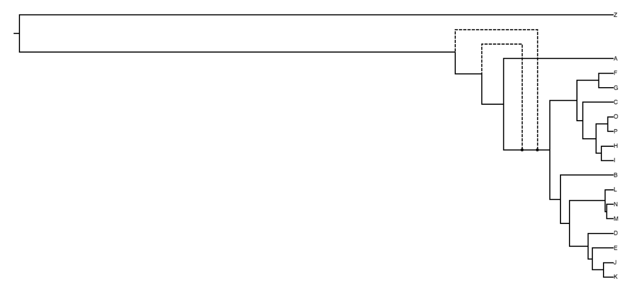 |
| 2 reticulations. Difficulty: Middle. The dependency of 2 reticulations makes inference not easy. | 2 reticulations. Difficulty: Hard. The reticulations are deep and dependent. |
| 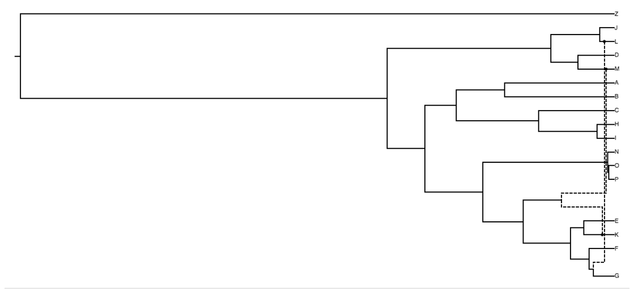 | 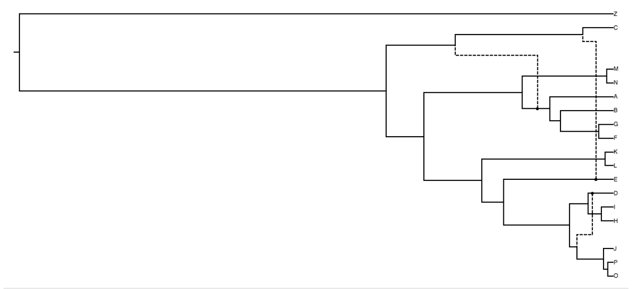 |
| 3 reticulations. Difficulty: Easy. | 3 reticulations. Difficulty: Middle. The dependency of up 2 reticulations makes inference not easy. |
| 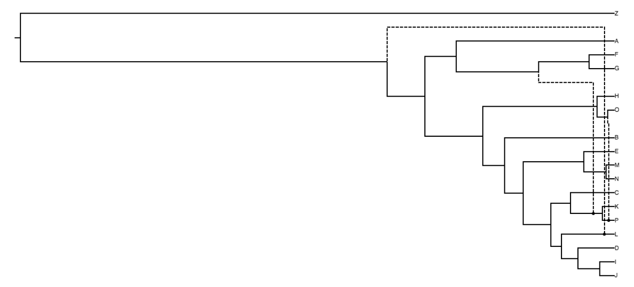 | 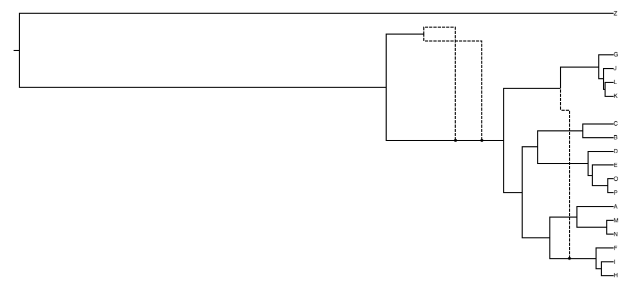 |
| 3 reticulations. Difficulty: Middle. The dependency of 2 reticulations above K and P makes inference not easy. | 3 reticulations. Difficulty: Hard. The reticulations are deep and dependent. |

| | |
|---|---|
| 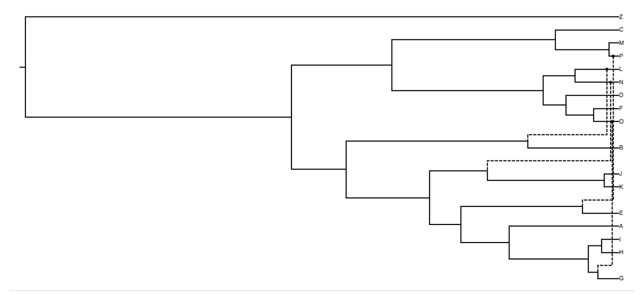 | 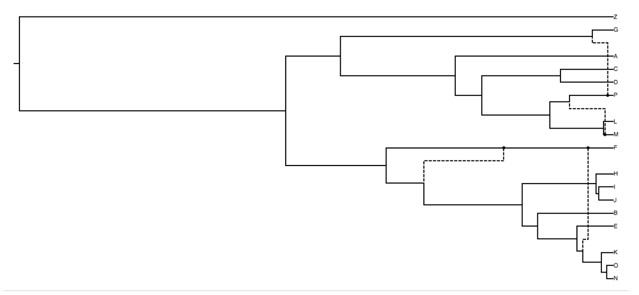 |
| 4 reticulations. Difficulty: Middle. The number of reticulations makes inference not easy. | 4 reticulations. Difficulty: Hard. The number of reticulations and dependencies make inference hard. |
| 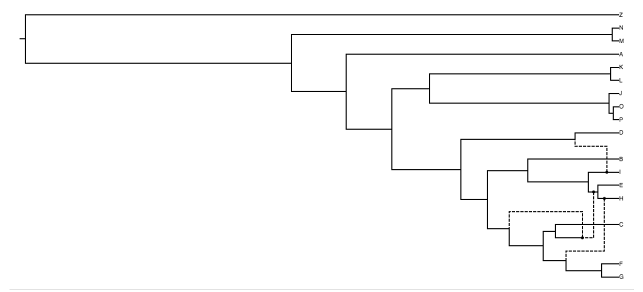 | 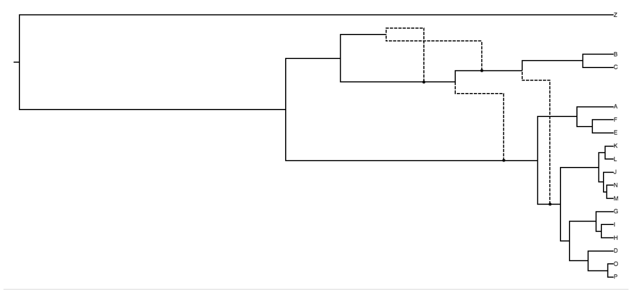 |
| 4 reticulations. Difficulty: Hard. The number of reticulations and dependencies make inference hard. | 4 reticulations. Difficulty: Hard. The number of reticulations and dependencies make inference hard. |
| 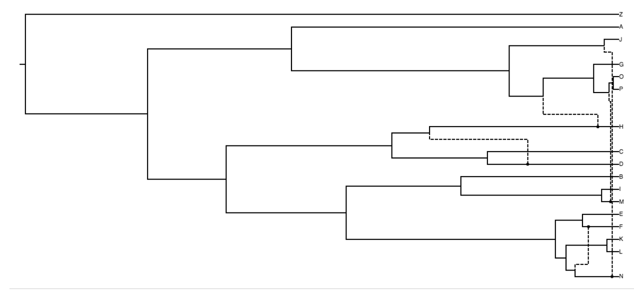 | 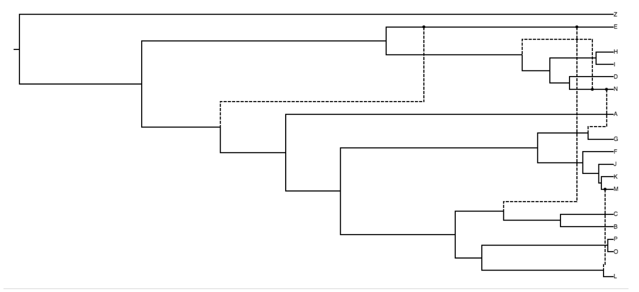 |
| 5 reticulations. Difficulty: Hard. The number of reticulations and dependencies make inference hard. | 5 reticulations. Difficulty: Hard. The number of reticulations and dependencies make inference hard. |
| 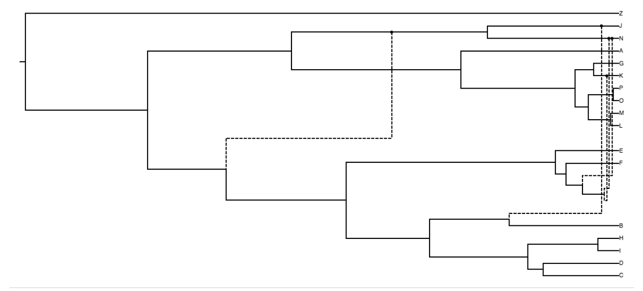 | 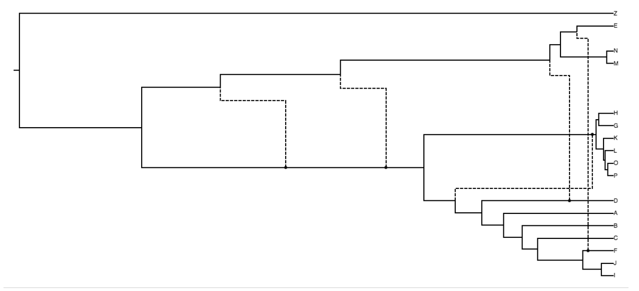 |
| 5 reticulations. Difficulty: Hard. The number of reticulations and dependencies make inference hard. | 5 reticulations. Difficulty: Hard. The number of reticulations and dependencies make inference hard. |

### 4 An example of the merger algorithm

Here we give a simple example to illustrate the merger algorithm. Fig. S1 shows the true network with 5 taxa as well as its all $\binom{5}{3} = 10$ subnetworks with 3 taxa. The subnetworks in Fig. S1(B) are obtained by restricting $\Psi$ to each combination of 3 taxa. Suppose the input sequence alignments are restricted to 10 combinations of 3 taxa, and inference algorithm is called. The inferred subnetworks are shown in Fig. S1. The inferred subnetworks are not necessarily identical to true subnetworks due to inference errors. Note that reticulation edges are missing in inferred $\Psi_1$, $\Psi_5$ and $\Psi_8$.

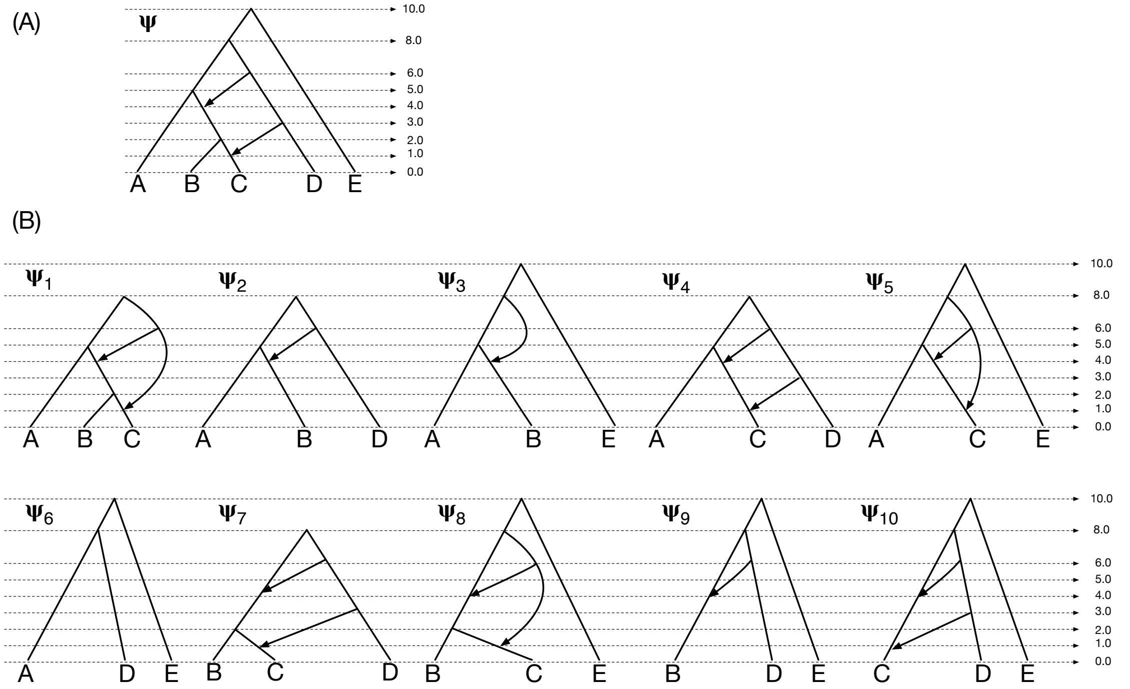


Figure S1: **The true network and its subnetworks.** (A) The true network $\Psi$, whose height of each node is indicated by the ticks. (B) The true subnetworks, whose height of each node is indicated by the ticks.

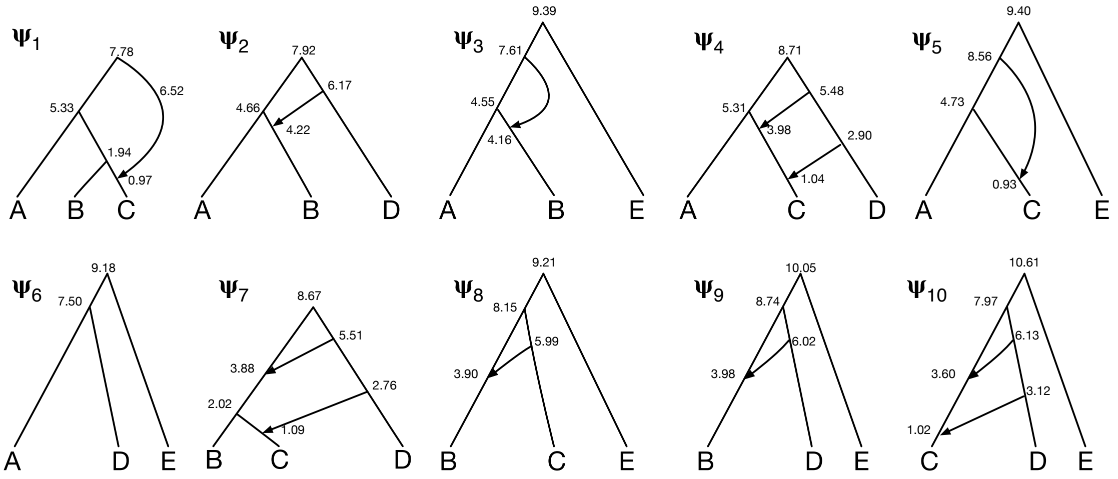


Figure S2: **The input to the merger algorithm.** Values near nodes of subnetworks are the height of each node. Colored dots indicate the corresponding nodes belong to the same disjoint set.

#### 4.1 Reconciling and summarizing the subnetworks

The first step is to reconcile the heights of nodes in each subnetworks. The disjoint sets of nodes are generated by mapping nodes in common binets in the subnetworks. Then the height of each node is assigned according to the average height of nodes in the same set. Fig. S3 shows the heights of nodes after reconciliation. Some disjoint sets are shown for illustration.

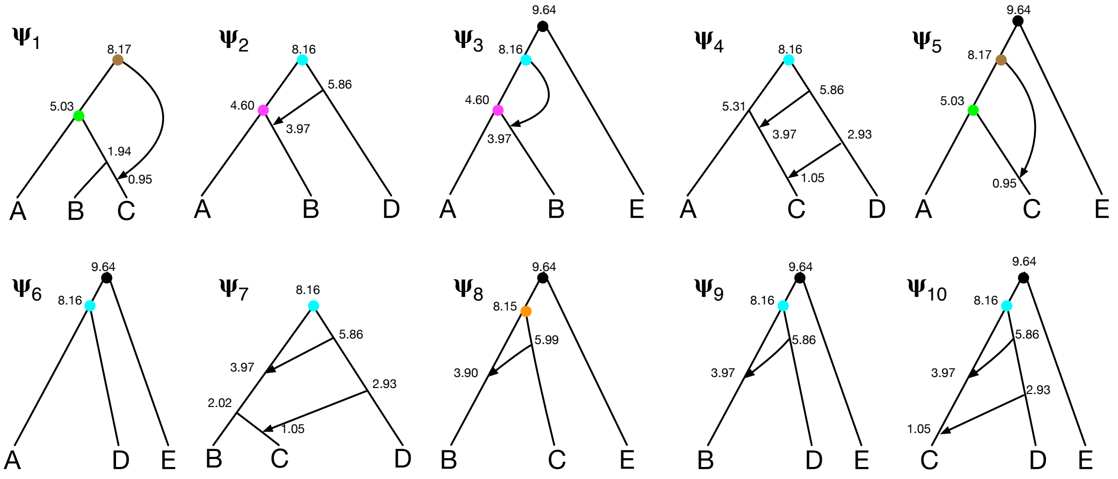


Figure S3: **The subnetworks after reconciling their heights.** Values near nodes of subnetworks are the height of each node.

Then, EHM is computed for every subnetwork.

$$\mathscr{M}_{\Psi_1} = \begin{array}{c} \\ \\ \\ \end{array}\begin{array}{ccc} A & B & C \\ \end{array}$$

$$\mathscr{M}_{\Psi_1} = \begin{pmatrix} - & [5.03] & [5.03, 8.17] \\ [5.03] & - & [1.94, 8.17] \\ [5.03, 8.17] & [1.94, 8.17] & - \end{pmatrix} \begin{matrix} A \\ B \\ C \end{matrix} \quad (\text{columns: } A, B, C)$$

$$\mathscr{M}_{\Psi_2} = \begin{pmatrix} - & [4.60, 8.16] & [8.16] \\ [4.60, 8.16] & - & [5.86, 8.16] \\ [8.16] & [5.86, 8.16] & - \end{pmatrix} \begin{matrix} A \\ B \\ D \end{matrix} \quad (\text{columns: } A, B, D)$$

$$\mathscr{M}_{\Psi_3} = \begin{pmatrix} - & [4.60, 8.16] & [9.64] \\ [4.60, 8.16] & - & [9.64] \\ [9.64] & [9.64] & - \end{pmatrix} \begin{matrix} A \\ B \\ E \end{matrix} \quad (\text{columns: } A, B, E)$$

$$\mathscr{M}_{\Psi_4} = \begin{pmatrix} - & [5.31, 8.16] & [8.16] \\ [5.31, 8.16] & - & [2.93, 5.86, 8.16] \\ [8.16] & [2.93, 5.86, 8.16] & - \end{pmatrix} \begin{matrix} A \\ C \\ D \end{matrix} \quad (\text{columns: } A, C, D)$$

$$\mathscr{M}_{\Psi_5} = \begin{pmatrix} - & [5.03, 8.17] & [9.64] \\ [5.03, 8.17] & - & [9.64] \\ [9.64] & [9.64] & - \end{pmatrix} \begin{matrix} A \\ C \\ E \end{matrix} \quad (\text{columns: } A, C, E)$$

$$\mathscr{M}_{\Psi_6} = \begin{pmatrix} - & [8.16] & [9.64] \\ [8.16] & - & [9.64] \\ [9.64] & [9.64] & - \end{pmatrix} \begin{matrix} A \\ D \\ E \end{matrix} \quad (\text{columns: } A, D, E)$$

$$\mathscr{M}_{\Psi_7} = \begin{pmatrix} - & [2.02, 5.86, 8.16] & [5.86, 8.16] \\ [2.02, 5.86, 8.16] & - & [2.93, 5.86, 8.16] \\ [5.86, 8.16] & [2.93, 5.86, 8.16] & - \end{pmatrix} \begin{matrix} B \\ C \\ D \end{matrix} \quad (\text{columns: } B, C, D)$$

$$\mathscr{M}_{\Psi_8} = \begin{pmatrix} - & [5.99, 8.15] & [9.64] \\ [5.99, 8.15] & - & [9.64] \\ [9.64] & [9.64] & - \end{pmatrix} \begin{matrix} B \\ C \\ E \end{matrix} \quad (\text{columns: } B, C, E)$$

$$\mathscr{M}_{\Psi_9} = \begin{array}{c} \\ \left( \begin{array}{lll} - & [5.86, 8.16] & [9.64] \\ [5.86, 8.16] & - & [9.64] \\ [9.64] & [9.64] & - \end{array} \right) \end{array} \begin{array}{l} \\ B \\ D \\ E \end{array}$$

with columns labelled $B$, $D$, $E$.

$$\mathscr{M}_{\Psi_{10}} = \left( \begin{array}{lll} - & [2.93, 5.86, 8.16] & [9.64] \\ [2.93, 5.86, 8.16] & - & [9.64] \\ [9.64] & [9.64] & - \end{array} \right) \begin{array}{l} C \\ D \\ E \end{array}$$

with columns labelled $C$, $D$, $E$.

Finally, an overall EHM $\mathscr{M}$ is computed. Take the entry $(B, C)$ for an example. The candidates for $(B, C)$ are $[5.03, 8.17]$, $[5.99, 8.15]$ and $[2.02, 5.86, 8.16]$. $[2.02, 5.86, 8.16]$ has the most elements, so it is selected as the entry in the overall EHM.

$$\mathscr{M} = \left( \begin{array}{lllll} - & [4.60, 8.16] & [5.03, 8.17] & [8.16] & [9.64] \\ [4.60, 8.16] & - & [2.02, 5.86, 8.16] & [5.86, 8.16] & [9.64] \\ [5.03, 8.17] & [2.02, 5.86, 8.16] & - & [2.93, 5.86, 8.16] & [9.64] \\ [8.16] & [5.86, 8.16] & [2.93, 5.86, 8.16] & - & [9.64] \\ [9.64] & [9.64] & [9.64] & [9.64] & - \end{array} \right) \begin{array}{l} A \\ B \\ C \\ D \\ E \end{array}$$

with columns labelled $A$, $B$, $C$, $D$, $E$.

#### 4.2 Generating a starting network and an order for leaf addition

The "outgroup" is set to E. The score of a subnetwork $\Psi_i$ is computed by $s(\Psi_i)+\sum_{1\le j\le k} d(\Psi_i, \Psi_j)$.

| $i$ | 1 | 2 | 3 | 4 | 5 | 6 | 7 | 8 | 9 | 10 |
|---|---|---|---|---|---|---|---|---|---|---|
| $s(\Psi_i)$ 1 | 1 | 1 | 1 | 1 | 1 | 0 | 1 | 1 | 1 | 1 |
| $\sum_{1\le j\le k} d(\Psi_i, \Psi_j)$ | 12 | 2 | 2 | 6 | 6 | 0 | 6 | 10 | 0 | 6 |

The subnetwork $\Psi_6$ with taxa A, D, and E, shown in Fig. S4(A) is selected as the starting network because it has the lowest backbone score. Then we need to generate an order of attaching new leaves. The guide graph whose nodes are the taxa set, and edges are computed according to reticulations in the subnetworks. Fig. S4(B) shows the guide graph. Topological sorting yields $[B, C]$ after removing A, D and E.

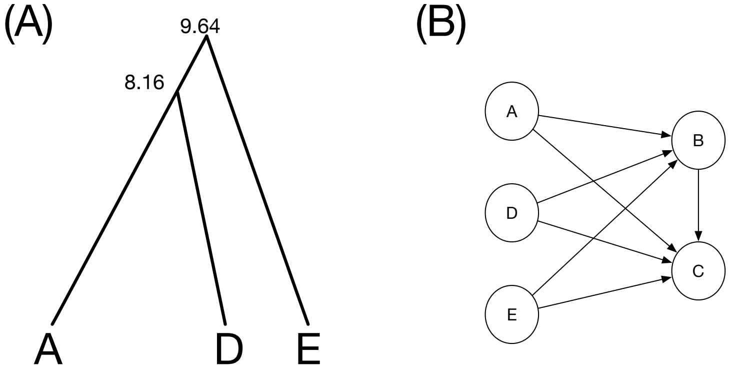


Figure S4: **A starting network and a guide graph for leaf addition.** (A) The starting network, and its heights of nodes are represented by the nearby values. (B) The guide graph whose nodes are represented by circles.

#### 4.3 Adding B

Attachments of B is extracted from subnetworks, as shown in Fig. S5. Attachments are clustered by the number of blue nodes in them. Therefore there are two clusters: $\{at_{\Psi_1}(B), at_{\Psi_7}(B)\}$ and $\{at_{\Psi_2}(B), at_{\Psi_3}(B), at_{\Psi_8}(B), at_{\Psi_9}(B)\}$. For the first cluster, $at_{\Psi_1}(B)$ is chosen (the two items are equivalent, so either one can be chosen). For the second cluster, $at_{\Psi_8}(B)$ is chosen, because the parent of B has the lowest height in it. Therefore, $H(B) = \{at_{\Psi_1}(B), at_{\Psi_8}(B)\}$.

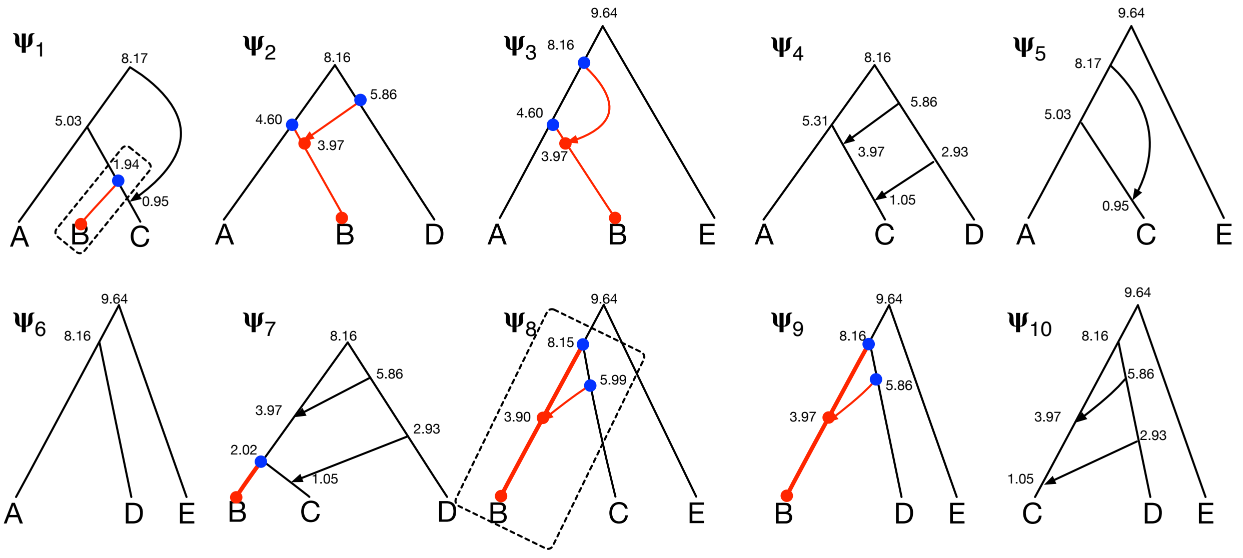


Figure S5: **Attachments of B.** Colored nodes and edges belong to an attachment of B in a subnetwork. Blue nodes are in set $rt(B)$ of corresponding subnetwork, and red nodes are in set $it(B)$ of corresponding subnetwork. Attachments surrounded by dotted rectangles are selected in $H(B)$.

Alg. 1 is called and sorted HT pairs in $\mathscr{H}$ are

- (4.60, A)
- (5.86, D)
- (8.16, A)
- (8.16, D)
- (9.64, E)

**Enumerate** tries to connect attachments in $\mathscr{H}$ with the draft network.

1. $at_{\Psi_1}(B)$ is tried. $(4.60, A)$ is resolved, and this yields the network in Fig. S6(A).
2. $at_{\Psi_8}(B)$ is tried. In the either permutation, after $(4.60, A)$ is resolved, the draft network is in Fig. S6(B). The node with in-degree 0 is removed and this yields the network in Fig. S6(A). Then $(5.86, D)$ and naturally all remaining HT pairs are resolved. This yields network in Fig. S6(C). The score of the network in Fig. S6(A) is 11, and the score of the network in Fig. S6(C) is 1, therefore the network in Fig. S6(C) is selected, and after reconciling its heights of nodes, we get a new backbone network in Fig. S6(D).

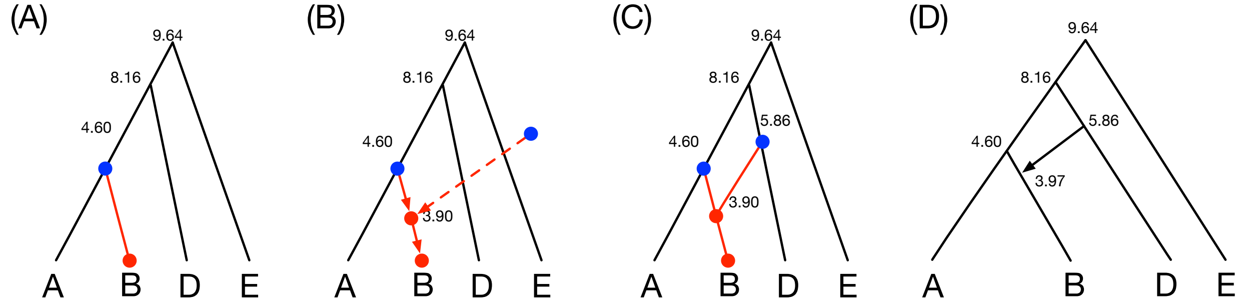


Figure S6: **Enumeration for B.** (A) The stored network when $(4.60, A)$ is resolved. (B) The draft network when examining $at_{\Psi_8}(B)$ with $(4.60, A)$ resolved. The arrows indicate the direction of branches in the attachment. The dotted branch is removed when storing this network. (C) The stored network when all HT pairs are resolved. (D) The new backbone network after adding taxon B and reconciling heights of nodes.

#### 4.4 Adding C

Attachments of C is extracted from subnetworks, as shown in Fig. S7. Attachments are clustered by the number of blue nodes in them. There are three clusters: $\{at_{\Psi_8}(C)\}$, $\{at_{\Psi_1}(C), at_{\Psi_5}(C), at_{\Psi_7}(C)\}$ and $\{at_{\Psi_4}(C), at_{\Psi_{10}}(C)\}$. For the first cluster, $at_{\Psi_8}(B)$ is chosen. For the second cluster, $at_{\Psi_5}(C)$ is chosen (the three items are equivalent, so either one can be chosen). For the third cluster, $at_{\Psi_{10}}(C)$ is chosen (the two items are equivalent, so either one can be chosen). Therefore, $H(C) = \{at_{\Psi_8}(C), at_{\Psi_5}(C), at_{\Psi_{10}}(C)\}$.

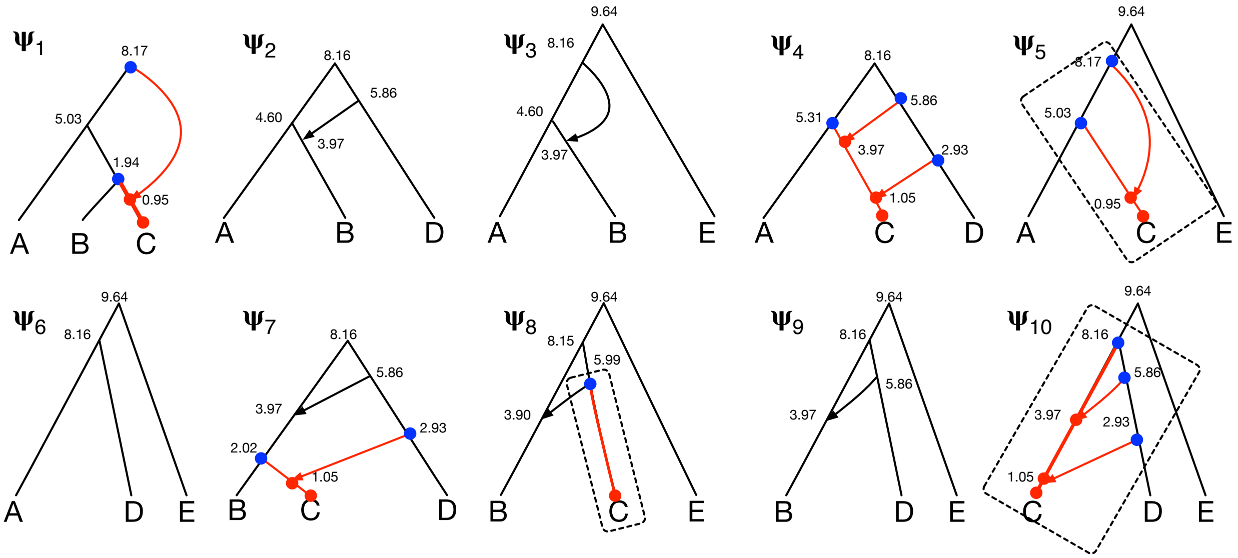


Figure S7: **Attachments of C.** Colored nodes and edges belong to an attachment of C in a subnetwork. Blue nodes are in set $rt(C)$ of corresponding subnetwork, and red nodes are in set $it(C)$ of corresponding subnetwork. Attachments surrounded by dotted rectangles are selected in $H(C)$.

Alg. 1 is called and sorted HT pairs in $\mathscr{H}$ are

- (2.02, B)
- (2.93, D)
- (5.03, A)
- (5.86, B)
- (5.86, D)
- (8.16, B)
- (8.16, D)
- (8.17, A)

- (9.64, E)

**Enumerate** tries to connect attachments in $\mathscr{H}$ with the draft network.

1. $at_{\Psi_8}(C)$ is tried. $(2.02, B)$ is resolved, and this yields the network in Fig. S8(A).
2. $at_{\Psi_5}(C)$ is tried. In the either permutation, $(2.02, B)$ is resolved, then $(2.93, D)$. This yields both networks in Fig. S8(A)(B).
3. $at_{\Psi_{10}}(C)$ is tried. Take one permutation in Fig. S8(C) for an example. $(2.02, B)$ is resolved, and yields Fig. S8(A). Then $(2.93, D)$ is resolved and draft network is in Fig. S8(D), note that the node with height 3.97 is removed when storing the draft network, so it yields the network in Fig. S8(F). When resolving $(5.03, A)$, the draft network is in Fig. S8(E), and the node with height 3.97 creates negative branch length thereby the draft network is discarded.

Note that the topology of the networks in Fig. S8(B)(F) are identical, the only difference is the height of one reticulation node. The score of the network in Fig. S8(A) is 19, while the score of the network in Fig. S8(B)(F) is 4, therefore the network in Fig. S6(B)(F) is selected, and after reconciling the heights of nodes, we get a new backbone network in Fig. S6(G).

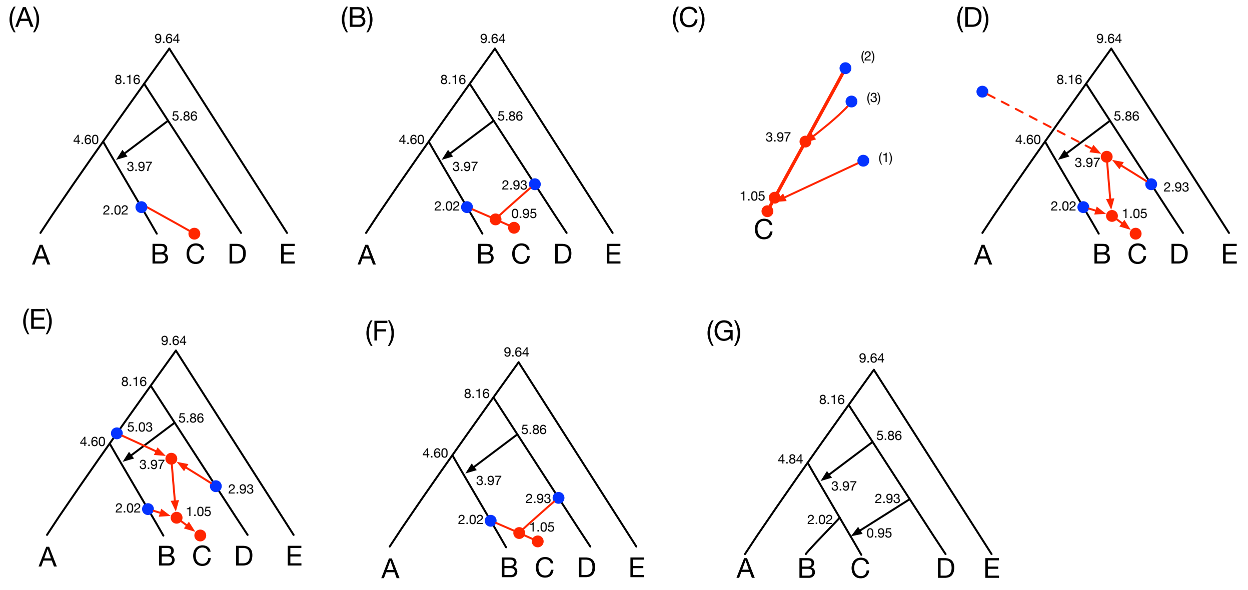


Figure S8: **Enumeration for C.** (A) The stored network when $(4.60, A)$ is resolved. (B) The stored network when $(4.60, A)$ and $(2.93, D)$ are resolved. (C) The attachment $at_{\Psi_{10}}(C)$ and the permutation of $rt_{\Psi_{10}}(C)$ with indices shown in parentheses. (D)(E) Draft networks when trying $at_{\Psi_{10}}(C)$. (F) stored network when trying $at_{\Psi_{10}}(C)$. (G) The final output.

Since all taxon are in the network in Fig. S6(G), it is the final output of merger algorithm. Its topology of is identical to the true network, and the heights of its nodes are close to the true ones.
